# CLT-seq as a universal homopolymer-sequencing concept reveals poly(A)-tail-tuned ncRNA regulation

**DOI:** 10.1101/2022.09.11.507502

**Authors:** Qiang Su, Yi Long, Jun Wang, Deming Gou

## Abstract

Dynamic tuning of the poly(A) tail is a crucial mechanism for controlling translation and stability of eukaryotic mRNA. Achieving a comprehensive understanding of how this regulation occurs requires unbiased abundance quantification of poly(A)-tail transcripts and simple poly(A)-length measurement using high-throughput sequencing platforms. Current methods have limitations due to complicated setups and elaborate library preparation plans. To address this, we introduce Central Limit Theorem (CLT)-managed RNA-seq (CLT-seq), a simple and straightforward homopolymer-sequencing method. In CLT-seq, an anchor-free oligo(dT) primer rapidly binds to and unbinds from anywhere along the poly(A) tail string, leading to position-directed reverse transcription with equal probability. The CLT mechanism enables the synthesized poly(T) lengths, which correspond to the templated segment of the poly(A) tail, to distribute normally. Based on a well-fitted pseudogaussian-derived poly(A)-poly(T) conversion model, the actual poly(A)-tail profile is reconstructed from the acquired poly(T)-length profile through matrix operations. CLT-seq follows a simple procedure without requiring RNA-related pre-treatment, enrichment, or selection, and the CLT-shortened poly(T) stretches are more compatible with existing sequencing platforms. This proof-of-concept approach facilitates direct homopolymer base-calling and features unbiased RNA-seq. Therefore, CLT-seq provides unbiased, robust, and cost-efficient transcriptome-wide poly(A)-tail profiling. We demonstrate that CLT-seq on the most common Illumina platform delivers reliable poly(A)-tail profiling at a transcriptome-wide scale in human cellular contexts. We find that the poly(A)-tail-tuned ncRNA regulation undergoes a dynamic, complex process similar to mRNA regulation. Overall, CLT-seq offers a simplified, effective, and economical approach to investigate poly(A)-tail regulation, with potential implications for understanding gene expression and identifying therapeutic targets.

## INTRODUCTION

Dynamic poly(A)-tail tuning is a critical mechanism that controls both mRNA translation and stability in all eukaryotic cells.^1, 2^ The importance of the poly(A) tail in biological processes is reflected in its occurrence in non-coding RNAs (ncRNAs) as well,^3, 4^ yet the regulation of poly(A)-tail-tuned ncRNA remains largely unknown.^5^ One reason for this limited understanding is the difficulty in base-calling homopolymeric sequences.^6^ However, creative approaches have been developed to address this issue. For instance, TAIL-seq^7^ and PAL-seq^8^ are pioneering methods that enable transcriptome-wide poly(A)-tail profiling on high-throughput sequencing platforms. The Tailseeker program, which uses machine learning, was applied in TAIL-seq for base-calling A-residues in homopolymer rows.^7, 9, 10^ Similarly, the internal standard and calibration curve method used in PAL-seq can estimate poly(A)-tail length.^8, 11^ The recent advancements in long-read sequencing by Nanopore^12, 13^ and PacBio^14-17^ have also improved the ability to detect homopolymeric sequences through repeated error correction.

These methods have expanded our understanding of the dynamic nature of the mRNA poly(A) tail, but their complicated setups limit their applicability to various scenarios. For instance, the Tailseeker 1/2 programs are only applicable to the Illumina 4-channel system, instead of a 2-channel system even with more efficient sequencing. Moreover, the Nanopore and PacBio-seq methods provide only the poly(A)-tail length of genes expressed at medium to high levels due to their low sequencing throughput.^18, 19^ While these methods have expanded our understanding of the dynamic nature of the mRNA poly(A) tail, their complicated setups limit their applicability to various scenarios. For example, the Tailseeker 1/2 programs are only applicable to the Illumina 4-channel system, despite the fact that there are more efficient sequencing options available with a 2-channel system. Additionally, the Nanopore and PacBio-seq methods provide only the poly(A)-tail length of genes expressed at medium to high levels due to their low sequencing throughput.

We introduce CLT-seq, a novel RNA sequencing technique that utilizes the Central Limit Theorem to overcome challenges in homopolymer sequencing. This approach involves synthesizing poly(T) sequences of varying lengths to enable straightforward measuring of the poly(A) tail. By randomly extending an oligo(dT) primer along the poly(A) tail, the resulting distribution of poly(T) lengths corresponding to the length-fixed poly(A) tails follows a normal distribution. From the obtained poly(T) length profile, we can reconstruct the poly(A) tail profile using a well-fitted pseudogaussian-derived poly(A)-poly(T) conversion model through matrix operations. However, stochastic phasing may result in overestimation of some homopolymer lengths, particularly for longer strings, reducing accuracy for individual poly(A) tails. To address this issue, we employ a pseudogaussian function that accounts for skewness and kurtosis in the poly(T)-length distribution converted from a single narrow peak of poly(A) tails. Additionally, CLT-seq reduces reverse-transcription biases for accurate transcript-abundance estimation. We validated this technique using mutual corroboration and derivation, simulation-measurement comparisons, Ligation-Mediated Poly(A) Tail (LM-PAT) Assays, and statistical analyses. Applying CLT-seq to common Illumina sequencing platforms, we comprehensively profiled the poly(A) tail of mitochondrial transcriptomes and demonstrated that poly(A)-tail-tuned ncRNA regulation is as complex as mRNA regulation. Overall, CLT-seq offers a simple and effective solution for unbiased RNA sequencing and poly(A) tail profiling.

## RESULTS

### Principle and feasibility of CLT-seq

We have developed CLT-seq, a technique for profiling poly(A) tails in multiple cellular contexts using a simple procedure without pre-treatment, enrichment, or selection (Fig. 1, a). The process involves an adapter-flanked anchor-free oligo(dT) randomly annealing to the poly(A) tail for the first cDNA synthesis. After excess primer depletion, the resulting cDNA couples with the second adapter through a splint-ligation approach and serves as a template for PCR-based library amplification.

**Fig. 1.**
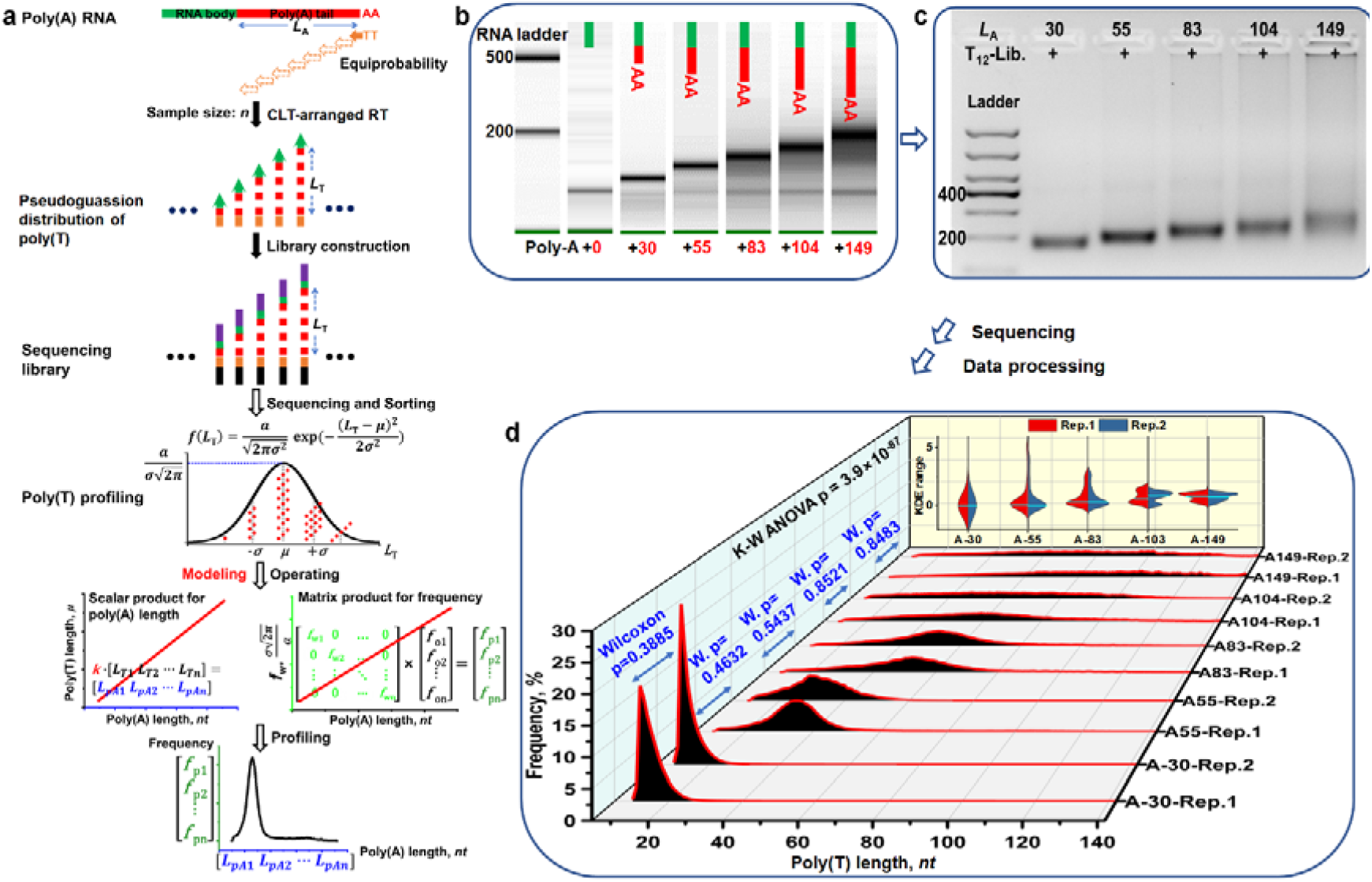
The principle and feasibility of CLT-seq on the Illumina platform. **a**) Experimental and computational outline of CLT-seq. **b**) Home-made spike-ins with varying poly(A)-tail lengths (red bar). The lengths are measured by Agilent 2100 bioanalyzer with RNA 6000 Pico Kit. **c**) Agarose-gel (3%) electrophoresis of the PCR-amplified libraries. **d**) 3-D plot CLT-arranged poly(T) length against counting frequency for a series of spike-ins with two replicates. Wilcoxon between A-30, A-55, A-83, A-104, and A-149 replicates p is 0.3885 (z=0.8624), 0.4632 (z=-0.7335), 0.5437 (z=0.6073), 0.8521 (z=0.1865), and 0.8483 (z=0.1913). Kruskal-Wallis ANOVA p is 3.9 × 10^−87^ (df=9). Splint violin with KDE range is displayed (median, cyan line).

The CLT mechanism enables the poly(A)-templated poly(T)-synthesis lengths to distribute normally, as the anchor-free oligo(dT) binds equiprobably to and rapidly unbinds from any complementary segment of the poly(A) tail until it is primed by a transcriptase. The number of randomly positioned oligo(dT)-binding events determines a sample size for a pseudogaussian distribution function (Function in Method), which converts poly(A) length into the CLT-managed poly(T) length. To model the pseudogaussian-based conversion, we generate a set of poly(A) tails through poly(A) polymerase-catalyzed spike-in RNA polyadenylation under time-controlled incubation. We then precisely measure the extended poly(A) lengths using an Agilent 2100 bioanalyzer (Fig. 1, b). The resulting poly(A)-tailed spike-ins and their corresponding first cDNAs, primed from non-anchored oligo(dT18) and anchored oligo(dT18VNN), are characterized by electrophoresis (Supplemental Fig. S1, a). The non-anchored oligo(dT) priming leads to diffuse first-cDNA components because it primes randomly along the poly(A) tail, compared to the anchored oligo(dT) priming at the fixed position on the poly(A)-tailed spike-in template. Libraries based on oligo(dT_12_) (Fig. 1, c), oligo(dT_18_), and oligo(dT_30_) (Supplemental Fig. S1, b) display a poly(A)-tail-dependent library-shifting tendency, but with different diffusion for the long poly(A) tails, attesting CLT’s controlled mechanism. Replicated experiments for a series of spike-ins show good reliability for poly(T) profiling in the Wilcoxon rank-sum test (Fig. 1, d). As the poly(A)-tail length increases, the poly(T) profile shifts to the right along with curve flattening in the poly(T)-length spectrum, which is confirmed by the Kruskal-Wallis ANOVA. The validation tests confirm the reliability of the CLT-seq technique for unbiased RNA sequencing and poly(A) tail profiling. In sequencing-based homopolymer-length determination, a wide distribution of homopolymer strings with a fixed length is usually observed instead of a single narrow peak.^7^ This results in a skewed length distribution due to the tendency for homopolymer lengths to be overestimated. Ideally, the mean length in the sequencing-length distribution should be equal to the actual homopolymer size. To measure synthesized length-fixed DNA homopolymers (A_20_, A_40_, and A_60_) stretches, we used the two-channel NovaSeq instrument which has a lower error rate and less variation than most other Illumina platforms.^20, 21^ For A_20_ and A_40_, the pre-defined poly(A) length is exactly equal to the mean value in the well-profiled length distribution (Supplemental Fig. S2, a and b). However, even though the mode value is fixed at 60 nt, a skewed length distribution due to homopolymeric length overestimation is generated from A_60_ (Supplemental Fig. S2, c). The read quality decreases as the homopolymer base is called one by one because of the stochastic phasing that often occurs at every ∼15 sequencing cycles.^22, 23^ The stochastic phasing recurs throughout the sequencing process, leading to sequencing-length overestimation. The standard base-calling algorithm adequately corrects this error at an early stage of base-calling. Although the Tailseeker base-calling software can significantly improve homopolymer-sequencing accuracy through an advanced base-correcting algorithm, a few instances of length-overestimating stretches are still observed in the 64-nt homopolymer sequencing, and overestimation is more noticeable in the 118-nt length determination. Despite these overestimations, a reliable measurement of poly(A)-tail length (up to 300 nt) is still achieved at genome-wide scale.^7^ We also provided a good reproducibility on CLT-seq poly(A)-length (up to 142 nt) measurements in our biological replicates (Fig. 1, d and Supplemental Fig. S3).

The CLT mechanism facilitates the distribution of CLT-shortened homopolymer lengths to follow a normal pattern. To demonstrate this, we have examined the sequencing-poly(T)-length distribution of various poly(A) tails ranging from A_30_ to A_149_, as shown in Supplemental Fig. S3. Even in the case of the longest tail, A_149_, we still observed a characteristic mode length of 80-nt (73 and 87 nt) using a pseudogaussian function fitting, which was found to be very close to the theoretical length of 75 nt. Despite the cumulative overestimation that may distort the sequencing-length distribution and make it appear flatter,^7^ our proposed pseudogaussian function provides a way to account for the skewness and kurtosis effects when modeling such data. Although Illumina’s standard base-calling algorithm has limited accuracy for measuring the various lengths of mammalian poly(A) tails,^5^ it is adequate for profiling CLT-shortened homopolymeric stretches for characterization purposes. Each poly(A) tail associated with a specific mRNA species contributes to a unique spectral signature characterized by its poly(A)-length wide distribution. The CLT mechanism ensures that every poly(A) component derived from these gene-specified transcripts undergoes reverse transcription while being subjected to CLT management. This process transforms the gene-specific poly(A)-distribution spectrum into a distinct poly(T) profile. In the following section, we will develop a conversion model between the mode poly(A) length and mode poly(T) length.

### Mathematical consideration for CLT-seq

The oligo(dT) consisting only of thymine can anneal to its complementary poly(A) segment, but it has a low binding affinity for reverse transcription (RT) at 37□.^24^ The poly(A) tail has a fixed number of binding sites for oligo(dT) (Fig. 2, a). When a binding event occurs, an oligo(dT)-poly(A) monovalent complex forms. This complex can either instantly unbind without transcriptase-loading assistance or become progressively more stable as the loading transcriptase extends the oligo(dT). Each binding event assigns a potential starting priming position on the poly(A) template for transcription. Assuming that binding events are randomly packed to define the sample size underlying the Central Limit Theorem (CLT) mechanism (Function in Methods), packed numbers representing binding positions on the poly(A) tail are averaged to determine the coincidentally oligo(dT)-priming position (Fig. 2, a, bottom). From this starting position, the poly(T) stretch synthesis is directed by the poly(A)-tail template. In probability theory, the length distribution of the synthesized poly(T) approximates a normal distribution as the sample size grows larger. Thus, the distribution of poly(T) length was simulated as the sample size ranged from 1 to 30 for T_12_ and A_55_ RT conditions (Fig. 2, b). The resulting profile was compared with an experimental conducting profile obtained from the T_12_ primed-spike-in-A_55_ CLT-seq. The two profiles showed strong similarity (Fig. 2, c), suggesting that the assumption of the CLT mechanism arranging length distribution is valid.

**Fig. 2.**
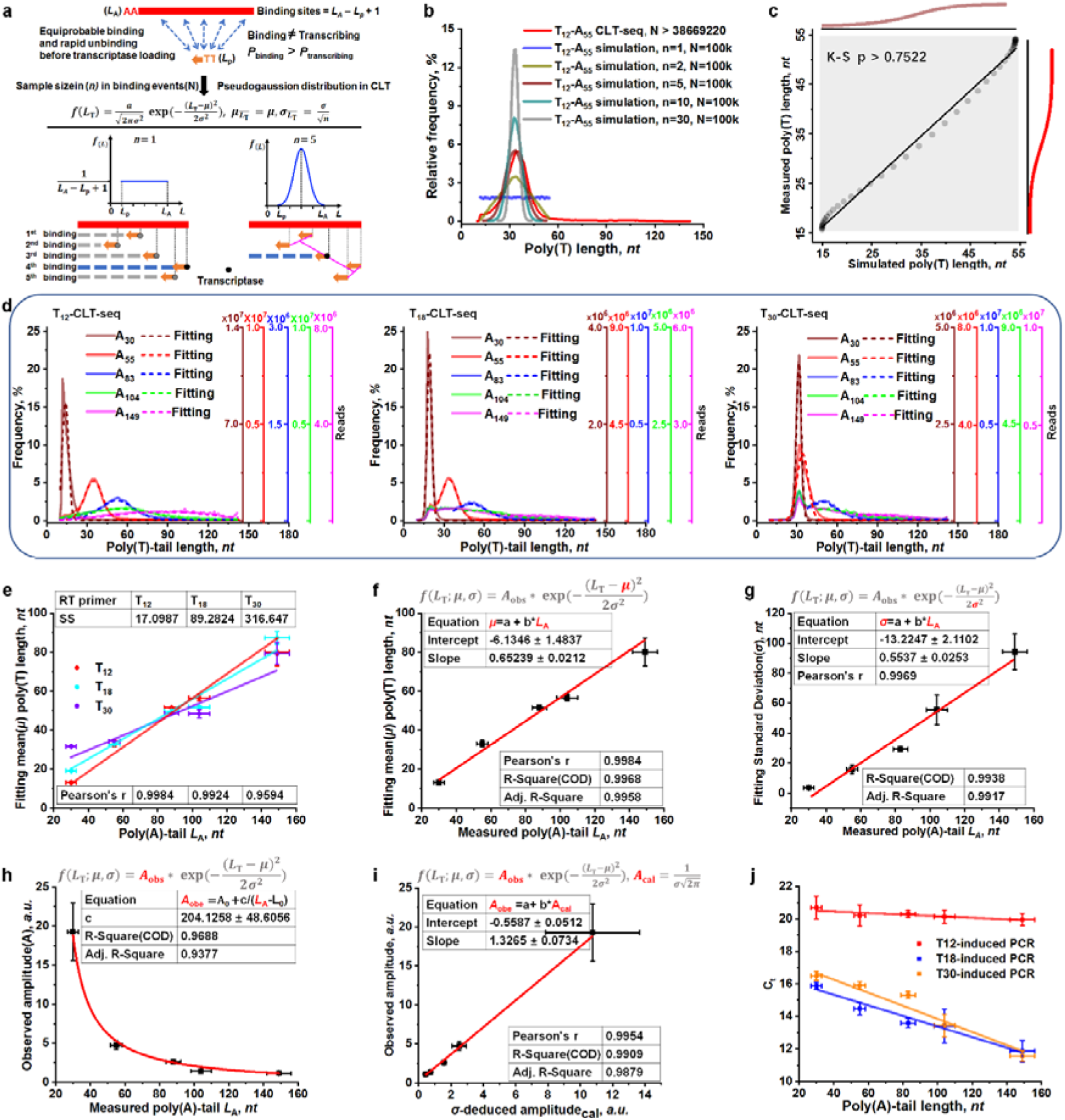
Mathematical consideration of CLT-seq. **a**) Theoretical interpretation of CLT-arranged poly(T)-length distribution as sampling size increases. The poly(A)-oligo(dT) complexes are more than the poly(A)-oligo(dT)-transcriptase complex. The oligo(dT) primer equiprobably bound to and instantly unbound from the poly(A) segment till it was primed by the loading transcriptase. The sample size (n) determines the number of the poly(T)-length-assigned binding events (first to fifth, for instance) that are randomly packed. The mean length corresponding to the *n* independent binding events (*n*=5, for instance) is close to the deserved length of the middle oligo(dT) (3^rd^). **b**) CLT-seq simulation under T_12_ and A_55_-fixed conditions and an experimental T_12_-primed-A_55_-template profile. **c**) The PP plot of poly(T) length ranging from 12 to 55 nt and Kolmogorov-Smirnov test (K-S test) for\ simulation-measurement comparison. K-S p is 0.7523 (z=0.6794). **d**) The overlapped poly(T)-length profiles generated from T_12_, T_18_, and T_30_-based CLT-seq. Counting frequency and reads are plotted versus the poly(T) length. **e**) T_12_, T_18_, and T_30_-based regression lines of poly(A) tail length versus fitting poly(T) mean length and fitted standard deviation. **f-g**) The correlation analysis between the measured poly(A)-tail length and output mean poly(T) length or standard deviation. Data are represented as mean ± s.e.m. for two biological replicates. **h**) The regression curve of poly(A) tail length (*L*_A_) versus frequency amplitude (A) from the T_12_-primed dataset. **i**) The correlation analysis between the standard deviation ()-deduced amplitude versus the observed amplitude in experimental poly(T)-length profiles in the T_12_-primed dataset. **j**) qPCR assessment of poly(A) tail-length preference corresponding to T_12_, T_18_, and T_30_ primers.

The T_12_-, T_18_-, and T_30_-primed poly(T) profiles (Fig. 2, d) for a range of spike-ins are overlaid and fitted with a pseudogaussian function (Supplemental Fig. S3 and S4). This confirms that there is a linear relationship between the input poly(a) length and the output mode poly(t) length or standard deviation (Fig. 2, e). Comparing the standard least square-fitting results in T12, T18, and T30-based datasets (Supplemental Fig. S5), we observed that the T12-based CLT-seq provides the most accurate model for poly(A)-poly(T) conversion (Fig. 2, f and g As the poly(A) tail length increases, the observed frequency amplitude decreases hyperbolically (Fig. 2, h). We have derived a hyperbolic equation (Equation in Methods) from the pseudogaussian function to predict the frequency amplitude from the fitted standard deviation. The measured amplitude and predicted value match well (Fig. 2, i), demonstrating the validity of the pseudogaussian function for interpreting the CLT mechanism. Additionally, the T12-based CLT-seq can effectively minimize bias induced by poly(A) tail length (Fig. 2, j), owing to the weak binding affinity of T_12_. The bias arising from the T_12_-primed PCR amplification efficiency may be overlooked with an approximately two-fold increase in yield, even over a tail length of 129 nt. However, there is a significant increase in T_18_ and T_30_-primed amplification efficiency with increasing tail length. This bias is likely present in anchor-free oligo(dT)-based sequencing, which is currently one of the most commonly used protocols^25^ for whole transcriptome profiling but has not been previously reported.

### Practical utility of CLT-seq

In CLT-seq, profiling the poly(A) tail involves assigning sequencing-determined relative abundance to sorted lengths. However, this approach may be susceptible to PCR bias. To investigate this potential bias, we analyzed the pattern of insert length (Supplemental Fig. S6) and poly(T)-length distribution (Fig. 3, a) at the different phases of PCR amplification. We conducted CLT-seq with T_12_ primer using a spike-in-A_83_, and directly amplified resultant cDNA by qPCR for real-time ΔR_n_ output. Our findings indicate that PCR amplification at different phases does not significantly alter the poly(T)/insert-length proportion across a wide GC-content range. This can be attributed primarily to the high processivity and exceptional fidelity of KAPA HiFi polymerase.^26^ While library construction requires appropriate PCR amplification, it is crucial to avoid over-amplification, particularly for rare cells or clinical samples. Since the CLT mechanism operates in various cellular contexts, we also investigated the effect of the oligo(dT)-primer/transcript-object ratio on the poly(T)-length distribution (Fig. 3, b). We subjected a spike-in-A55 to transcription using T_12_ primer-transcript ratio ranging from 0.2:1 to 200:1 with ten-fold serial dilutions. Subsequently, we quantified the produced cDNA by qPCR. Even over a 3-order-of-magnitude ratio range, the poly(T)-distribution spectrum shifted by at most two bases. A previous study yielded similar results, indicating that increasing the primer/transcript ratio in the RT system had almost no impact on the poly(T) profiling.^27^

**Fig. 3.**
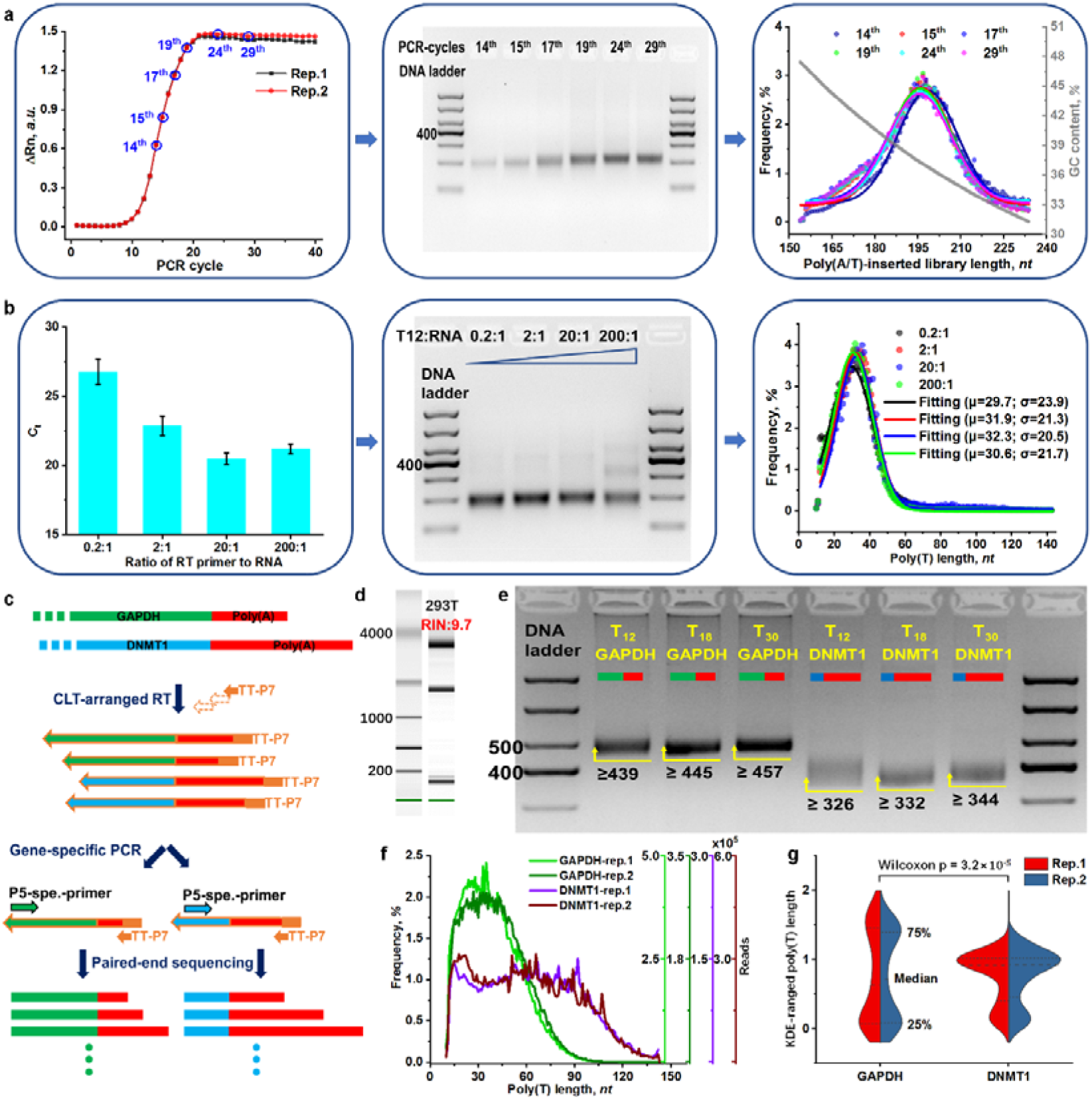
Practical utility of CLT-seq. **a**) Poly(T) profiling at the different phases of PCR amplification. The PCR cycle number of 14^th^, 15^th^, 17^th^, 19^th^, 24^th^, and 29^th^ in two replicates are marked by circles (left), and the cleaned-up libraries are identified by electrophoresis with 3% agarose gel (middle). The poly(T)-length profile of each library is overplotted versus counting frequency and GC content (right). Arrows represent the stepwise procedure. **b**) Primer-transcript ratio in poly(T) profiling. The different ratio-induced 1^st^ cDNA is quantified by qPCR with relative C_t_ value (n=3, triplicates) (left), and the cleaned-up libraries are characterized by electrophoresis with 3% agarose gel (middle). The pseudogaussian fitted poly(T) profiles are overlapped with a mean () and standard deviation () (right). **c**) The stepwise procedure of gene-specific poly(T) length determination by CLT-seq. **d**) An integer total RNA isolated from HEK293T. **e**) Characterizing the libraries of T_12_, T_18_, and T_30_-based CLT-seq by electrophoresis. **f**) The poly(T) length of GAPDH and DNMT1 plotted versus the counting frequency and sequencing reads (n=2, two replicates). **g**) A split violin based on Kernel density estimation (n=2, two replicates). The dash and dot lines are median and quantiles, respectively.

The reliability and utility of CLT-seq in complex environments were evaluated by conducting T_12_, T_18_, and T_30_-directed CLT-seq on the total RNA of HEK293T using gene-specific PCR amplification (Fig. 3, c and d). The cDNA generated by CLT was directly amplified using human GAPDH and DNMT1 primers, and the desired libraries were identified through gel electrophoresis (Fig. 3, e). Based on T_12_, T_18_, and T_30_, the minimum size of the PCR amplicon for the GAPDH library was found to be 439, 445 (439+6), and 457 (439+18) nt, respectively. Meanwhile, based on the same oligo(dT) primers, the minimum size of the amplicon for the DNMT1 library was found to be 326, 332 (326+6), and 344 (326+18) nt, respectively. DNMT1’s library was more diffuse than GAPDH, indicating that the mean poly(A)-tail length of DNMT1 is longer than that of GAPDH according to the diffusion ranging function (Function in Methods). This finding is supported by the comparison of the CLT-seq sequencing poly(T)-length distribution (Fig. 3, f). Further confirmation of a significant difference between DNMT1 and GAPDH was obtained through the Wilcoxon-Mann-Whitney test with p = 3.2 × 10^−5^ (z=-4.16), as revealed in Fig. 3, g. These results demonstrate the reliability and utility of the CLT-mechanism operation in multiple cellular contexts.

### Transcriptome-wide profiling poly(A) tail by CLT-seq

We conducted a transcriptome-wide profiling of poly(A) tails using a standard procedure (Fig. 4, a). However, we encountered read-length limitations and overlap in Illumina paired-end sequencing that made it necessary to ensure that the poly(T) length reading from the R2 end was not shorter than the poly(A) length reading from the R1 end (Fig. 4, a, bottom library). We observed that the poly(T)-length distribution obtained from R2 exhibited a right-shift profile compared to the poly(A)-length distribution obtained from R1 in the GAPDH and ZNF544 (Supplemental Fig. S7). This indicates that the paired-end R_1_ and R_2_ must be in the correct position to accurately map genes and output poly(A) sequences, respectively. To evaluate transcriptome-wide mapping performance, we used two types of oligo(dT) primers in RNA-seq experiments. We found that an anchor-free and short T_12_ primer yielded a comparable pattern of read-mapping distribution (Fig. 4, b) to typical representatives of anchored^28, 29^ and non-anchored^25, 30^ long-oligo(dT) primers, highlighting the pronounced oligo(dT)-specific poly(A)-dependent reverse transcription. The mapping coverage over each transcript illustrated the 3’-end preference of transcripts confirmed by the R1-based mapping result across six primer-based datasets.^31^ The paired R2 carries more poly(T) sequence and less mRNA information, making it unsuitable for gene-specific identification and applicable only to outputting poly(T) sequence. To obtain the poly(A)-tail-length frequency, we used the scalar-product formula (Fig. 4, c; Formula in Methods) and initially rescaled the poly(T) length obtained from the well-fitted conversion to the poly(A) length. However, we observed that the pseudogaussian-shaped poly(T) profile flattened as the poly(A)-tail length increased (Fig. 2, d), leading to a reduced frequency (amplitude) that had to be offset by a weighted frequency accounting for length overestimation (Formula in Methods). We obtained the poly(A)-tail-length frequency by taking the matrix product of the length-weighting factor and the poly(T)-length-sorted frequency (Fig. 4, c; Formula in Methods). Furthermore, we computed the mean poly(A)-tail length by taking the matrix product of the poly(A)-tail length and the weighted frequency (Formula in Methods).

**Fig. 4.**
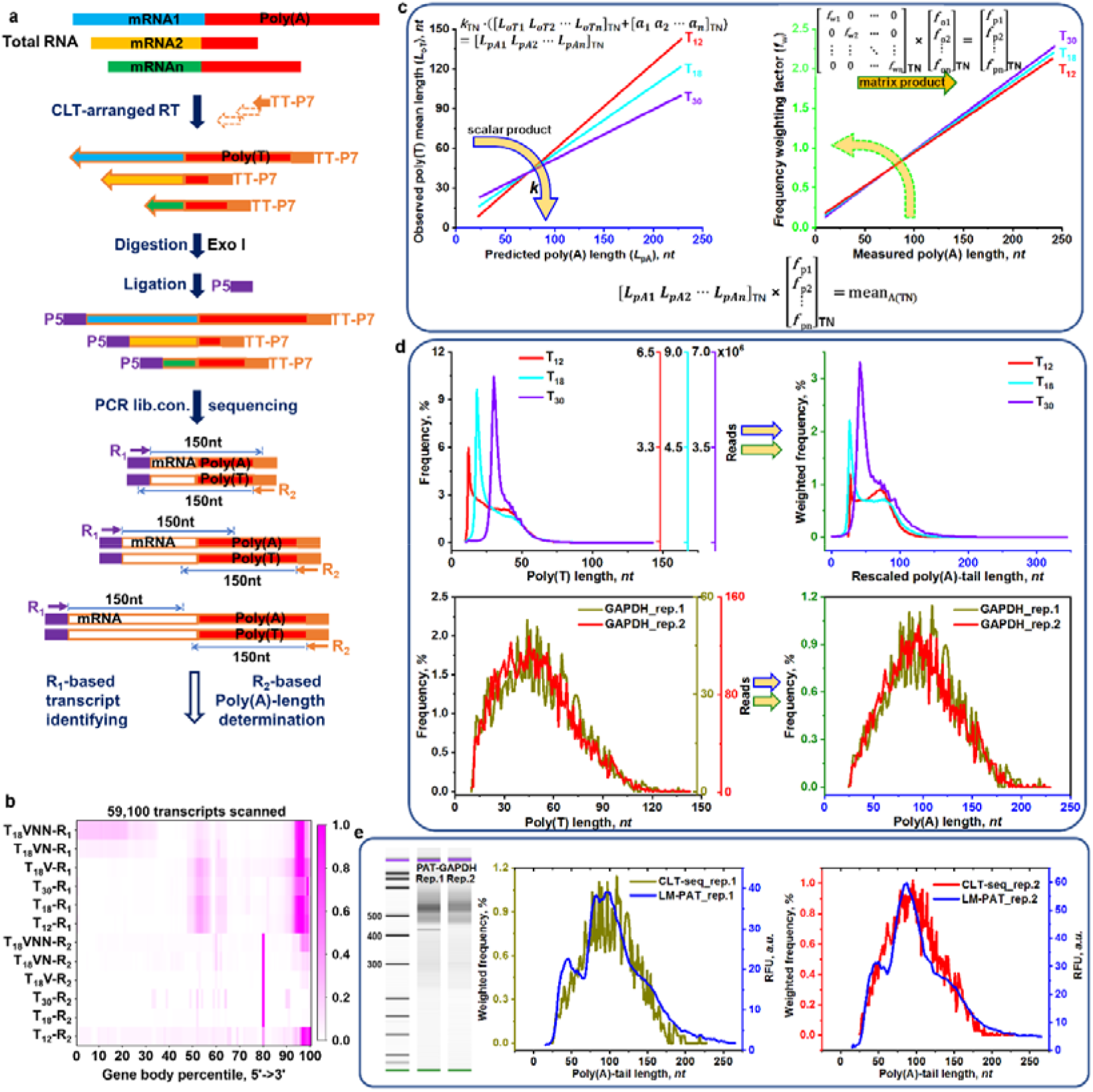
Transcriptome-wide poly(A)-tail length determination and validation. **a**) The stepwise procedure of transcriptome-wide tail-length determination by CLT-seq. **b**) Mapping read coverage over 59 100 transcripts across six oligo(dT) primers based on paired-end R_1_ and R_2_ reads. **c**) Scalar and matrix operation for poly(A) rescaling, frequency weighting, and mean poly(A)-tail length determining. T_12_ (red), T_18_ (cyan), or T_30_ (violet) primer takes a different set of parameters. The arrow means the operating process. **d**) Global profile of poly(A) tail in HEK293T with T_12_ (red), T_18_ (cyan), or T_30_ (violet)-based CLT-seq. GAPDH-specific poly(A) tail profile from T_12_-based CLT-seq, n=2 (two replicates). **e**) Poly(A) length validation by LM-PAT assay for GAPDH with two replicates. PAT assay product identified by Agilent 2100 bioanalyzer with DNA chip in left. PAT assay profile (blue) with RFU (*a*.*u*.) overlapped the poly(A) length distribution (red) with weighted frequency (%) restored from CLT-seq data in the right panel.

In this study, global poly(A)-tail profiles were compared across T12, T18, and T30-primed poly(T) datasets (Fig. 4, d, up). The analysis revealed similar poly(A)-tail-length distribution among the three datasets, with two peaks observed at approximately 25-nt and 88-nt poly(A) lengths in the T_12_ and T_18_ datasets. Notably, the length of 88-nt highlights the estimated mean global poly(A) length of 88.4 nt in HEK293T by PAL-seq.^8^ Since the CLT-seq followed an unbiased procedure without pre-treatment, enrichment, and selection of the total RNA, the naturally high abundance^9^ of the short-tailed transcripts is likely to be profiled by the CLT-seq and usually missed in traditional library-preparing procedures like oligo(dT)-bead based capturing. To test the reliability of the CLT-seq in multi-transcript poly(A)-tail profiling, a good correlation was observed between the biological replicates of GAPDH measurement (Fig. 4, d, down). However, some degree of fluctuation in frequency was noted, which appeared to correlate with sequencing depth (the total number of gene-identified reads) (Supplemental Fig. S8). A minimum of ∼100 reads was necessary to obtain a clear and smooth poly(a)-tail profile. In addition, we performed a ligation-mediated poly(A) tail assay (LM-PAT in Methods) with two replicates to further investigate the reliability of CLT-seq. The LM-PAT^32^ followed an enrichment- and pre-treatment-free procedure similar to CLT-seq, and the results showed agreement between CLT-seq and PCR-amplicon electrophoretic profiles of GAPDH (Fig. 4, e).

### Poly(A)-tail length control in ncRNA regulation

The mitochondrial transcriptome contains a wide variety of poly(A)-tailed mRNAs and ncRNAs, which makes it possible to sample full sets of cellular transcriptomes.^33^ The length of the poly(A) tail is known to be closely tied to translation efficiency during early embryonic development or the cell cycle,^8, 34^ making embryonic HEK293T cells and lung cancer A549 cells ideal for extensive poly(A)-tail studies using CLT-seq (Supplemental Fig. S1, c).We aligned the R1-decoded sequence to human sapiens mitochondrion (NC_012920.1) to identify all mitochondrial transcripts (37 RNA symbols and D-Loop), and then used the paired R2 reads to restore the poly(A)-tail profile (Fig. 5, a). In HEK293T and A549, we observed different patterns of mitochondrial-transcript abundance between intra-group and inter-group transcriptome (Fig. 5, b). Our measurements of mean tail length for literature-reported mitochondrial-mRNAs14 agreed with those obtained from CLT-seq.^14^

**Fig. 5.**
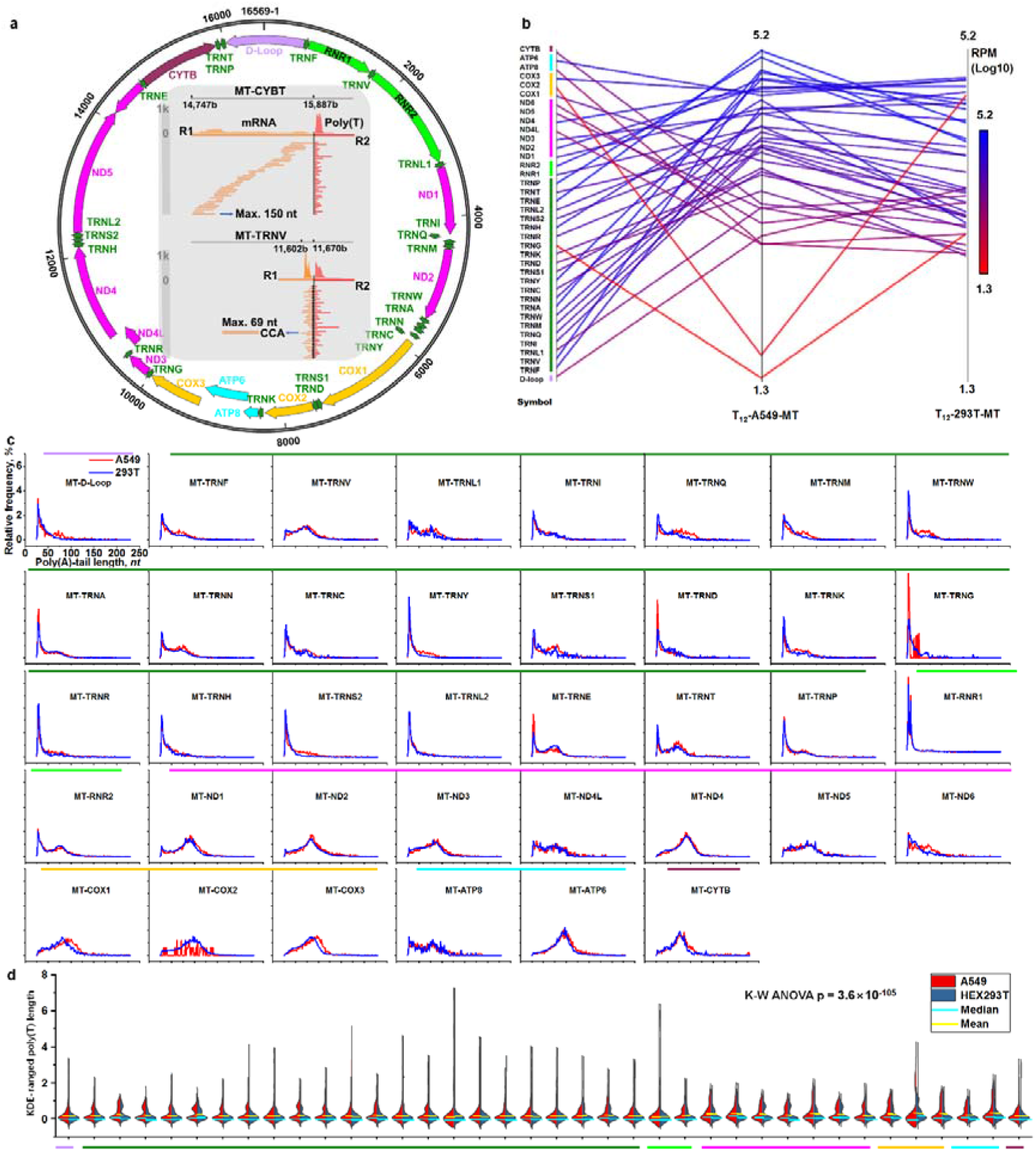
Poly(A)-tail length analysis of whole mitochondrial transcripts. **a**) Map of the human mitochondrial transcriptome with annotated gene features regarding NC_012920.1 (outer tracker) and two poly(A)-tail profiling strategies for individual transcript (inner). In an example of a lengthy transcript (mitochondrial-CYTB), the sequence with a maximum length of 150 nt from paired-end R_1_ is mapping onto 1141-nt CYTB mRNA reference, and the corresponding poly(T) length is outputting. For short transcript (mitochondrial-TRNV), the sequence from R_1_ directly aligns to the ncRNA-poly(A) tail junction for the poly(T) tail profiling. **b**) Transcript expression abundance. All transcripts are in type and subclass order with different colors. The abundance of RPM is plotted parallelly with the transcript symbol in the HEK293T and A549 dataset. **c**) The restored Poly(A)-tail profile by CLT-seq. The overlapped profiles of HEK293T and A549 tally counting frequency for each position with single-based resolution. **d**) The split violin plot of all mitochondrial transcripts with Kruskal-Wallis ANOVA analysis.

Profiling poly(A) tails by only using mean or median length is not sufficient to fully reflect the length of the poly(A) tail spectrum. Instead, we assigned sequencing-determined relative abundance to sorted lengths to create a profile for each mitochondrial poly(A) tail in HEK293T or A549 (Fig. 5, c). Despite having similar mean lengths, each mitochondrial poly(A)-tail profile was distinct. Some profiles (ND3 versus CYTB and ND6 versus RNR2) exhibited a bimodal length distribution, likely due to poly(A)-tail pruning during different phases of gene expression regulation.^35^ We observed a positively-skewed distribution in nonfunctional ncRNA (D-Loop), indicating poly(A)-tail-promoted degradation.^36^ To confirm these new findings, T_12_, T_18_, and T_30_-primed profiles overlapped for each mitochondrial transcript, exhibiting a similar tail-length pattern (Supplemental Fig. S9). We used the KDE method to normalize the poly(A)-tail profile with a Gaussian-shaped *Kernel* (Fig. 5, d). There is a significant difference between all gene-specific profiles. There was an overall increase in poly(A)-tail length in A549 tRNAs, potentially indicating their coupling with the specific upregulation of tRNA-derived small RNA (tsRNA) in lung cancer.^37, 38^ CLT-seq helped us understand the biogenesis and regulation mechanism of tsrnas.

To assess the reliability of discriminating subtle differences between profiles, we performed CLT-seq on two biological replicates of HEK293T (Supplemental Fig. S10) and A549 (Supplemental Fig. S11). There was good correlation between the two replicates across all mitochondrial transcripts. The correlation coefficient in ncRNAs with a short tail was higher than that of mRNAs with a longer poly(A) tail, most likely due to larger variation in long tail measurement. However, the long-length variation effect in CLT-seq only shifted the profile to some extent and did not change the fundamental pattern underlying the poly(A)-tail profile.

### Characterizing poly(A)-tail profile at transcriptome-wide scale

The Kruskal-Wallis ANOVA analysis showed a difference in mitochondrial poly(A)-tail profiles between HEK293T and A549, but the results were imprecise and unclear in many pairwise comparisons. Thus, to obtain a more precise statistical analysis, we utilized the Pearson correlation method for both the HEK293T (Fig. 6, a) and A549 (Fig. 6, b) datasets. By using this method, we were able to assign a more accurate Pearson’s correlation-coefficient value to each pairwise comparison. The resulting two-dimensional correlation pattern provides an exact quantitative interpretation of multiple endogenous poly(A) tails, even across different cell lines with varying characteristics.

**Fig. 6.**
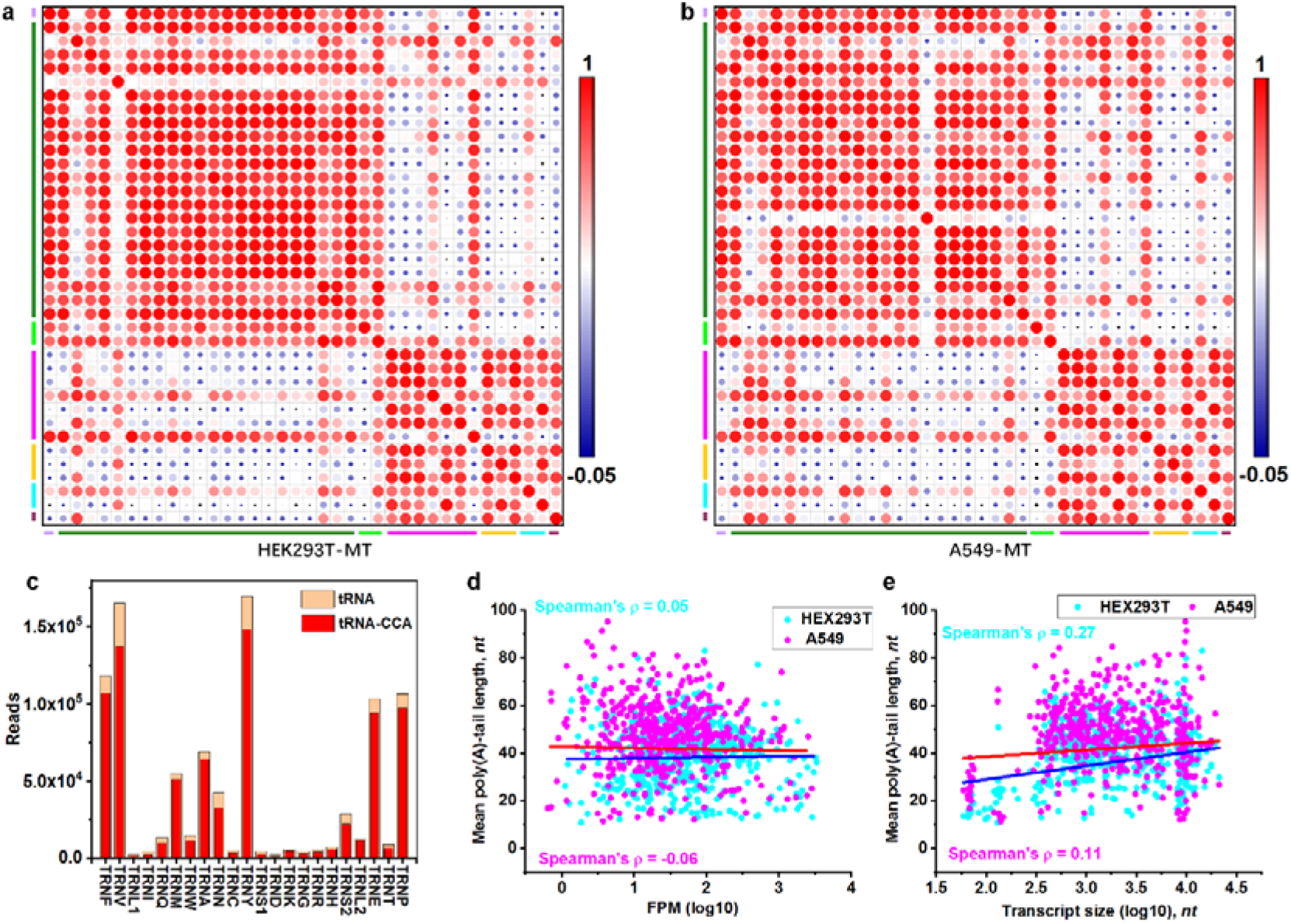
The featured poly(A)-tail. **a)** and **b**) Correlation scatter plot of pairwise comparison across all mitochondrial transcripts. Pearson correlation ecoefficiency (r) is related to the color and diameter of each spot in HEK293T and A549 datasets. **c**) The read count of CCA marked tRNA and all identified tRNA across all mitochondrial tRNAs in HEK293T. **d)** and **e**) Correlation of mean tail length with transcript abundance and size in HEK293T (cyan) or A549 (magenta) cells (n=1, Spearman correlation). Comparison of poly(A)-tailed ncRNA transcript abundance determined by T_12_-based CLT-seq between HEK293T and A549.

The poly(A) tail is commonly added to the 3’ CCA nucleotides at the 3’ end of the acceptor arm on all mitochondrial tRNAs in the T12-based (Fig. 6, c), T_18_-base (Supplemental Fig. S12, a), and T_30_-based dataset (Supplemental Fig. S12, b). On average, around 80% of all mitochondrial tRNAs are matured with the CCA tail (Supplemental Fig. S12, c). To confirm this high percentage, T_12_, T_18_, and T_30_-based CLT-seq were conducted in parallel, resulting in consistent accounting-for results across all mitochondrial tRNAs. Notably, a small proportion (about 0.3%) of mitochondrial tRNAs were found to have two CCA repeats (5’-NCCACCA-3’) at the 3’ end of the acceptor arm, indicating that these tRNAs had undergone degradation by the cytosolic tRNA nucleotidyltransferase 1 (TRNT1) system.^39^ This discovery provides strong evidence and confirms that TRNT1-mediated tRNA clearing is not limited to nuclear-coded tRNAs but also extends to mitochondrial-coded tRNAs. This is the first time that the 3’-end CCACCA (two repeats) sequences at mitochondrial tRNA have been directly observed, supporting the notion that the TRNT1-based tRNA clearing system functions in both cytosolic and mitochondrial contexts.

In our analysis of T12-based data from HEK293T and A549, we determined the mean tail length of transcripts with low abundance (Fig. 6, d). Interestingly, we found that the mean tail length did not correlate with RNA abundance but rather with transcript size (Fig. 6, e). This finding is consistent with previous observations regarding the relationship between median tail length and 3’-UTR length. Additionally, CLT-seq allows for the characterization of the poly(A) tail of all types of APA-spliced isoforms. To illustrate this capability, we plotted an isoform-specific poly(A)-tail profile at each APA site on the same 3’-UTR across different genes (Supplemental Fig. S13). We identified unique isoforms by assembling specific 3’-end UTR and poly(A) tail sequences for each type. Furthermore, we were able to profile the poly(A) tails of cross-intron spliced isoforms in individual genes. These isoforms contain skipped regions that allow for differentiation between the same APA site on the 3’-UTRs (Supplemental Fig. S14).

## DISCUSSION

The poly(A) tail plays a crucial role in eukaryotic RNA quality control,^40^ yet transcriptome-wide analyses of poly(A)-tail length have only become possible recently with the development of TAIL-seq^7^ and PAL-seq^8^ methods that overcame technical limitations in homopolymer base-calling. In recent years the Nanopore technology^12^ and PacBio system^14-16^ also proved to overcome homopolymer-reading barriers. However, these sequencing methods require complex library preparation workflows that can introduce biases. Additionally, there are still challenges in applying these methods for homopolymer-reading. To address these issues, we developed CLT-seq, which uses a simple workflow and straightforward base-calling approach based on a universal homopolymer-sequencing concept. By avoiding enriching and selecting steps, CLT-seq reduces most of artificial biases. The non-anchored oligo(dT) priming mechanism circumvents GC bias, while the short oligo(dT) helps mitigate tail-length bias. The randomized splint format used in CLT-seq also reduces bias during the introduction of the second adaptor.^41^ Furthermore, CLT-arranged transcription requires only low-input total RNA (_≤_ 500 ng), and the buffer contains divalent cations strong enough to shear long transcripts into proper fragments for short-read Illumina sequencing.^42^ The result is an unbiased RNA-seq at the transcriptome-wide scale achieved through these concerted efforts.

During traditional library preparation procedures and oligo(dT) primer-based RT, the short oligo(dT) binds and unbinds from its target segment of the poly(A) tail until primed by coincidentally-loading transcriptase. At this point, the CLT mechanism arranges the poly(A)-template-directed poly(T)-synthesis length to distribute normally, resulting in a gene-specific poly(A)-length distribution range that shrinks to approximately half of its original size, facilitating easy homopolymer sequencing. However, it is important to note that the length of some homopolymer strings may be overestimated due to stochastic phasing, particularly for long homopolymers. In addition to measuring A30 to A149 tails, we also measured an A∼198 tail using the CLT-seq method. However, the poly(T)-length profile induced by the A∼198 tail could not be explained with a pseudogaussian function at all. This may be due to excessive length-cumulative overestimation, which obscures the pseudogaussian spectrum of the poly(T)-length distribution and prevents setting of pseudogaussian parameters for modeling. To reliably and reasonably estimate poly(A) length, we recommend using CLT-seq to measure mean/mode/median poly(A) lengths below ∼150 nt. The well-fitted pseudogaussian-derived model underlying the weighted KDE method can appropriately interconvert between the poly(A) length (<149 nt) distribution and poly(T) length distribution, neutralizing overestimation effects. Since most human mRNA median poly(A)-tail lengths fall within the range of 50 to 100 nt,^8^ the CLT-seq method can effectively characterize most poly(A) tails in this range. Moreover, the shortened homopolymer can be easily read by advanced sequencing technologies such as Tailseeker, PacBio, and Nanopore. As a result, the CLT model can serve as a universal concept for poly(A) tail profiling, particularly on the Illumina platform, which has widespread availability and a well-tested bioinformatic pipeline.

We utilized CLT-seq on the Illumina platform to analyze the global poly(A)-tail profile in HEK293T cells and found two distinct peaks at approximately 25 nt and 88 nt, respectively. While the ∼88 nt peak is consistent with previous measurements obtained via PAL-seq,^8^ the presence of the ∼25 nt peak has rarely been reported in literature. This could be attributed to a transcript selection bias that filters out short-tail transcripts during traditional library preparation procedures and oligo(dT) primer-based RT, which preferentially select for longer-tailed transcripts. Despite this technical limitation, we believe that there is likely a considerable abundance of short-tailed transcripts resulting from regular metabolism of long poly(A) tails, conservation of ∼25-30 nt footprint tails of highly expressed genes, and random RNA degradation.^9^ These transcripts cannot be accurately captured or quantified through current sequencing methods, highlighting the need for approaches such as CLT-seq that can overcome this bias and enable the profiling of short-tail transcripts. In addition to profiling global poly(A)-tail length in HEK293T cells, we also conducted the first comprehensive analysis of poly(A)-tail length in the human mitochondrial transcriptome using CLT-seq. We observed that mitochondrial ncRNAs possess various poly(A) tails, indicating that poly(A)-tail-stimulated ncRNA degradation is a complex process under coordinated control. We further investigated the relationship between poly(A)-tail profiles and gene expression dynamics in lung cancer cells. Compared to normal cells, we identified a bimodal profile in many mitochondrial tRNAs in lung cancer cells. This altered pattern based on the poly(A)-tail-profile correlation can reliably track gene expression dynamics and characterize rare events. Overall, CLT-seq provides a more comprehensive sampling of total RNA sets and sheds new light on the regulation and function of poly(A) tails in cells.

## METHODS

### Data analysis

#### 1. Data pre-processing

The published-software pipeline processes the pair-end raw data: FastQC (v.0.11.8) (Babraham Bioinformatics), cutadapt (v.2.10)^43^, trimmomatic (v.0.39)^44^, hisat2 (v.2.2.1)^45^, samtools (v.1.7)^46^, and RSeQC package^47, 48^.

#### 2. Poly(T) profiling

For paired-end R_1_, poly(A) calling is preceded by RNA body, while for R_2_, poly(T) calling precedes RNA-body. R_1_ as forward read primarily calling RNA body and R_2_ as reverse read primarily calling poly(T) stretch are deployed to identify gene-specific transcript and output its corresponding poly(T) length, respectively. We firstly identified genes from the trimmed R1 reads aligned to the reference hg19_RefSeq.bed using the FPKM_count.py script. The sequence at the end of 3’-UTR from the identified gene packs the gene-specific R1-read ID from massive pair-end reads, and then the R2-read with the same index-ID was pooled together for poly(T)-length outputting.

#### 3. The code for poly(T) profiling is as following

The **code** for gene-specific grouping from R1:

grep -B1 ’UTR_sequence’ trimmed_input_R1.fastq | sed

’s/1:N:0:ATCTCGTA+ATAGAGAG(barcode)//g’ | awk NR%3==1 > R1_ID.txt

The **code** for R2-read pooling with the outputted R1_ID:

for i in $(cat R1_ID.txt)

do

grep -A1 $i trimmed_input_R2.fastq >> R2_reads_indexed_ID.txt

done

The **code** for gene-specific grouping:

for ((i=10; i<144; i++))

do

egrep-c T{$i}> R2_reads_indexed_ID.txt >> abundance_assigned_sorted_tail-length.txt done

#### 4. Read-mapping distribution

The raw data is treated following pipeline to generate .bam file that is processed by read_distribution.py script with reference hg19_RefSeq.bed. The acquired tags per Kb are normalized by sequencing depth.

#### 5. Mapping coverage

The .bam file is treated by geneBody_coverage.py script with reference hg19_RefSeq.bed.

#### 6. Nucleotide frequency

The .bam file is processed by read_NVC.py script with reference hg19_RefSeq.bed.

#### 7. Expression abundance quantification

The .bam file is processed by FPKM_count.py script with reference hg19_RefSeq.bed.

### Mathematical operation

#### 1. Weighted KDE method

To transcriptome-widely obtain the poly(A)-tail profile, the acquired poly(T)-length distribution is processed *via* matrix operation based on this verified model using the kernel density estimation (KDE) method. In statistics, KDE with a gaussian kernel is often used to estimate the probability density function. Given CLT-based pseudogaussian kernel, a weighted KDE with function:

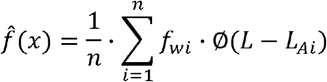

is applied to restore poly(A)-length profile (probability density function) where 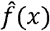 is the estimating probability density function, *f*_*wi*_ indicates the weighting factor for poly(A) length of i nt, and Ø (*L* − *L*_*Ai*_)is pseudogaussian function for i nt poly(A) length.

#### 2. Scalar product operation

The sorted poly(T) length is rescaled to the poly(A) tail length by a scalar product with function (Fig. 4, c, left):

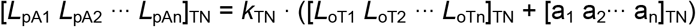

where *L*_oTn_ and *L*_pAn_ are the observed poly(T) length and the resulting poly(A) length in length order, respectively, _TN_ denotes the oligo(dT) primer (T_12_, T_18_ and T_30_), *k* is a scale factor, and a_n_ is a constant for all length n (a_1_=a_2_=⋯=a_n_).

#### 3. Matrix product operation

The counting frequency (density) of poly(A)-tail length (variable) is the matrix product of the indexed weighting factor and the observed frequency of the poly(T) length with the following function (Fig. 4, c, right):

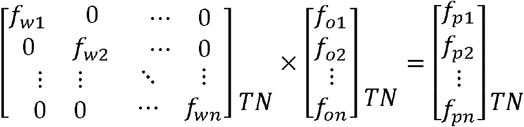

where *f*_wn_ in n×n matrix and *f*_on_ in n×1 matrix is the weighting factor and the observed frequency of poly(T) length of n nt, respectively. The weighted factors are listed orderly in a square diagonal matrix with poly(A)-length-dependent diagonal entries. This a weighted frequency should offset length-reduced frequency to reflect the transcript abundance-determined poly(A)-tail profile. Therefore, the weighted KDE method enables the CLT-based poly(A)-tail profile to approximate the actual tail-length distribution.

#### 4. Mean length (expected value)

The mean poly(A)-tail length is the matrix product of the poly(A) length and the weighted frequency. It follows that

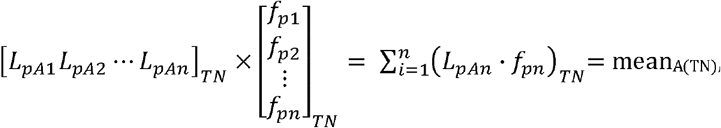

where *L*_pAn_ and *f*_pn_ represents the rescaled poly(A) length and the corresponding weighted frequency, respectively; mean_A(TN)_ denotes the mean poly(A) tail length (expected value) (Fig. 4, c, bottom).

### Pseudoguassian function

A pseudoguassian distribution function describes bell-like curve with 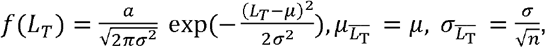 where *L*_T_ is the length of poly(T) in 1^st^ cDNA,*f*(L_T_) denotes the counting frequency (density), *μ* and *σ* is the population mean and standard deviation, respectively, a is constant, 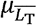 and 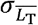 indicates the sampling mean and sampling standard deviation, respectively, and n is the sample size.

### Frequency amplitude-standard deviation Equation

According to the pseudogaussian function, a bell-like curve of CLT-based poly(T) length distribution flattened as rising poly(A)-tail length (Fig. 2d), leading to a reduced amplitude (counting frequency). The frequency amplitude is inversely proportional to the standard deviation (*σ*) as expressed by equation: 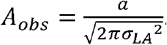.

#### Sequencing library diffusion range function

The library diffusion range (length-distribution range, R) is well-defined by an equation: R = *L*_A_-*L*_p_+1. The mean poly(A)-tail length of DNMT1 with strong diffusion is accordingly longer than that of GAPDH (*R*_DNMT1_ > *R*_GAPDH_ leading to *L*_A-DNMT1_ > *L*_A-GAPDH_ with fix-length primer (*L*_p_))

#### Cell culture

HEK293T cells were cultured in DMEM high glucose (catalog no. SH30022.01, HyClone) supplemented with 10% FBS (catalog no. 10100147, Thermo Fisher). In addition, A549 cells are cultured in F-12K (Gibco/Invitrogen) supplemented with 10% FBS. The cell lines grow at 37 _°_ and 5% CO_2_ with saturating humidity.

#### RNA isolation

All total RNAs were extracted from cells following the worksheet in RNAiso Plus (catalog no. 9109, TaKaRa Biotechnology) and resuspended at the appropriate concentration(s) in RNase-free water. The RNase-free water was used throughout the RNA-related assays. Total RNA was checked on 2100 Bioanalyzer RNA picochip and then aliquoted into 5 μg for long-term storing at -80 _°_.

#### Poly(A) DNA sequence

PCR with HIFI KAPA master mix (2×) according to the following mixture: 25 μl 2× HIFI KAPA master mix, 10 μl cDNA, 1 μl (50nM) synthesis DNA template,1 μl one of the two universal forward and reverse primers (10 μM), and H2O top up.

A20

5’-CACCTCTCTATACACTCTTTCCCTACACGACGCTCTTCCGATCTAAAAAAAAAAAAAAAAAAAAGATCGGAAGAGCACACGTCTGAACTCCAGTCACATCTCGTA-3’

A40

5’-CACCTCTCTATACACTCTTTCCCTACACGACGCTCTTCCGATCTAAAAAAAAAAAAAAAAAAAAAAAAAAAAAAAAAAAAAAAAGATCGGAAGAGCACACGTCTGAACTCCAGTCAC ATCTCGTA-3’

A60

5’-CACCTCTCTATACACTCTTTCCCTACACGACGCTCTTCCGATCTAAAAAAAAAAAAAAAAAAAAAAAAAAAAAAAAAAAAAAAAAAAAAAAAAAAAAAAAAAAAGATCGGAAGAGCACACGTCTGAACTCCAGTCACATCTCGTA-3’

#### Spike-in RNA and polyadenylation

0.5 μL of 20 μM spike-in RNA was added in 20 μL polyadenylation buffer with 0.4 μL of 5 U/μL Poly(A) Polymerase (part no. P7460L, QIAGEN) and 2 μL of 10 mM ATP for 0, 1-, 3-, 5-, and 10-min incubation at 37_°_:

Spike-in

5’-CACCUCUCUAUACACUCUUUCCCUACACGACGCUCUUCCGAUCUGAUCGTAAUGGCUUAG-3’

We noted the polyadenylated reaction started instantly (0 min incubating) after blending all reagents together.

#### Library preparation

500 ng of total RNA was subjected to 5 μl RT reaction with 0.5 μL of 1 μM primer T_12_, T_18_, T_30_, T_18_V, T_12_VN, and T_18_VNN:

T_12_:

5’-TACGAGATGTGACTGGAGTTCAGACGTGTGCTCTTCCGATCTAGCGTAGCTTTTTTTT TTTT-3’

TTTT-3′:

5’-TACGAGATGTGACTGGAGTTCAGACGTGTGCTCTTCCGATCTAGCGTAGCTTTTTTTT TTTTTTTTTT-3’

T_12_:

5′-TACGAGATGTGACTGGAGTTCAGACGTGTGCTCTTCCGATCTAGCGTAGCTTTTTTTT TTTTTTTTTT-3′

T_30_:

5’-TACGAGATGTGACTGGAGTTCAGACGTGTGCTCTTCCGATCTAGCGTAGCTTTTTTTTTTTTTTTTTTTTTTTTTTTTTT-3’

T_18_V:

5’-TACGAGATGTGACTGGAGTTCAGACGTGTGCTCTTCCGATCTAGCGTAGCTTTTTTTT TTTTTTTTTTV-3’

T_12_VN:

5’-TACGAGATGTGACTGGAGTTCAGACGTGTGCTCTTCCGATCTAGCGTAGCTTTTTTTT TTTTTTTTTTVN-3’

T_18_VNN:

5’-TACGAGATGTGACTGGAGTTCAGACGTGTGCTCTTCCGATCTAGCGTAGCTTTTTTTT TTTTTTTTTTVNN-3’

1 μl RT buffer mix (5×), 1 μl 200 U/μl of Super MMLV Reverse Transcriptase (catalog no. MD028, FAPON), 0.5 μl of 10 mM dNTP (each), and 3.3 mM Mg^2+^ at 37° C for 30 min. And then, 1 μL of 20 U/μl of Exonuclease I (catalog no. M0293S, New England Biolabs) was added directly to the RT mix for 30 min at 37°C and 20 min at 80°C. The resulting 1^st^ cDNA was mixed with 1 μL of 10 μM splint-second adaptor that was incubated at 95 °C for 5 min, slowly cooled to 25 °C:

5’-CACCTCTCTATACACTCTTTCCCTACACGACGCTCTTCCGATCTNNNNNN-invT-3’

5’-p-AGATCGGAAGAGCGTCGTGTAGGGAAAGAGTGTATAGAGAGGTG-invT-3’ in 20 μl ligation with 1 μl of 400 U/μl T4 DNA ligase (catalog no. M0202L, New England Biolabs) and 2 μl DNA ligase Buffer (10×). After 2 hours at 16°C, the final mix was then amplified by PCR with HIFI KAPA master mix (2×) according to the following mixture: 25 μl 2× HIFI KAPA master mix, 10 μl cDNA, 13 μl H_2_O and 1 μl one of the two universal forward and reverse primers (10 μM):

universal-forward-primer:

5’-AATGATACGGCGACCACCGAGATCTACACCTCTCTATACACTCT*T-3’

(*:phosphorothioate) universal-reverse-primer:

5’-CAAGCAGAAGACGGCATACGAGATGTGACTGGAGT*T-3’ (*:phosphorothioate)

The PCR reaction was then performed in a thermocycler as follows: 95 °C for 5 min, 23× (95 °C for 15 s and 60 °C for 30 s). The amplified library was then purified with 1.8× Ampure XP DNA Beads and resuspended in 20 μl H_2_O. Finally, one microliter of the library was checked by Qubit 4 Fluorometer with dsDNA HS (high sensitivity) Assay Kit (Invitrogen). Since there was not any clean-up step in the workflow, the library preparation in negative control yields a variable number of 4-types primers-introduced dimers or concatemers, which is minimized by the deserved target cDNA competition in the mRNA sample. Moreover, the non-specific amplicon without flow cell binding sites and sequencing primer binding sites are not likely to be sequenced.

#### Illumina sequencing

The purified PCR libraries were submitted to the Hoplox for Illumina sequencing (Novaseq 6000, two channel). The data output was according to a yield set.

#### RNA standards and Spike-ins

The poly(A) lengths were precisely measured by Agilent 2100 bioanalyzer as standard spike-ins carrying poly(A) tail of 30±2, 55±3, 83±4, 104±4, and 149±6 nt, respectively.

#### GAPDH and DNMT1 electrophoresis assay

The 500 ng of total RNA was subjected to a 5 μl RT reaction with 0.5 μL of 1 μM T_12_, T_18_, or T_30_, 1 μl RT buffer mix (5×), 0.5 μl of 10 mM dNTP (each), 1 μl of 200 U/μl Super MMLV Reverse Transcriptase (catalog no. MD028, FAPON), 3.3 mM Mg^2+^ at 37°C for 30 min. And then, 0.5 μL of 20 U/μl Exonuclease I (catalog no. M0293S, NEB) was added directly into the RT mix for 30 min at 37°C and 20 min at 80°C. Subsequently, the mix was then amplified by PCR with 25 μl HIFI KAPA master mix (2×), 1 μl universal reverse primer (10 μM) and 1 μl forward primer GAPDH-tag or DNMT1-tag (10 μM),

GAPDH-tag:

5’-CACCTCTCTATACACTCTTTCCCTACACGACGCTCTTCCGATCTCCTCAACGACCACTTTGTCAAG-3’

DNMT1-tag:

5’-CACCTCTCTATACACTCTTTCCCTACACGACGCTCTTCCGATCTCTGTTGTGTGAGGTTCGCTTATCAAC-3’

The PCR reaction was then performed in a thermocycler as follows: 95 °C for 5 min, 20× (95 °C for 15 s and 65 °C for 30 s).

#### LM-PAT assay

The 500 ng of total RNA was subjected to a 5 μl RT reaction with 0.5 μL of 100 nM the phosphorylated T_12-18_ and 1 μl RT buffer mix (5×) at 65 °C for 5 min, quickly cooled to 37 °C. 15 μl of the following prewarmed mixture is added and incubated at 37 °C for 30 min. For each reaction add: 4 μl RT buffer mix (5×), 2 μl DNA ligase Buffer (10×), 0.5 μl of 10 mM dNTP (each), 0.5 μl of 10 mM ATP, 1 μl of 200 U/μl Super MMLV Reverse Transcriptase (catalog no. MD028, FAPON), 1 μl of 400 U/μl T4 DNA ligase (catalog no. M0202L, New England Biolabs). At the end of the incubation, while still at 37 °C, 0.5 μL of 1 μM T_12_ was added and incubated at 16 °C for 2 hours. Subsequently, 1 μl of 200 U/μl Super MMLV Reverse Transcriptase (catalog no. MD028, FAPON) was added and incubated at 37 °C for 30 min and 65 °C for 30 min.

Finally, the cDNA was pre-amplified by PCR with 2 μl universal reverse primer (10 μM) and 2 μl forward primer GAPDH-1-tag (10 μM). The PCR reaction was performed in a thermocycler as follows: 95 °C for 5 min, 15× (95 °C for 15 s and 60 °C for 30 s). The pre-amplified product was further amplified with two universal primers for the 2100 Bioanalyzer and Illumina sequencer. The PCR reaction was performed in a thermocycler as follows: 95 °C for 5 min, 20× (95 °C for 15 s and 60 °C for 30 s).

## AVAILABILITY

The code used for data analysis is available at https://github.com/QiangSu/polyA_profiling. The code for poly(T) length determination is also available in methods.

## ACCESSION NUMBERS

All sequencing raw data and processing data have been deposited on NCBI GEO (https://www.ncbi.nlm.nih.gov/geo/) under the accession number GSE202256. Any other relevant data are available from the authors upon request.

## SUPPLEMENTARY DATA

Supplementary Data are available at online.

## ACKNOWLEDGEMENT

We thank Ms Qinyi Xia for providing HEK293T cells and Dr Kang Kang and Mr Haoyang Yin for providing A549 cells, Ms Shasha Zhang, and Mr Yongjie Liang for sequencing library preparation and preprocessing of sequencing data. We also thank all the members of the Prof Deming Gou laboratory for critical and useful discussions.

## FUNDING

We acknowledge financial support from the National Natural Science Foundation of China (82170070, 91739109, 81970053), Guangdong Provincial Key Laboratory of Regional Immunity and Diseases (2019B030301009), Shenzhen Municipal Research Grant (JCYJ20190808123219295, JCYJ20210324120206017) and Shenzhen-Hong Kong Collaborative Innovation Research (SGDX20201103095404019).

## CONFLICT OF INTEREST

The authors declare no financial interests.

## Notes

### Competing Interest Statement

The authors have declared no competing interest.

### Summary of Updates

The figures and dicriptions have been improved.

