## supplemental material for "CLT-seq as a universal homopolymer-sequencing concept reveals poly(A)-tail-tuned ncRNA regulation"

**This PDF file includes:**

Figures S1 to S14

**Fig. S1**.

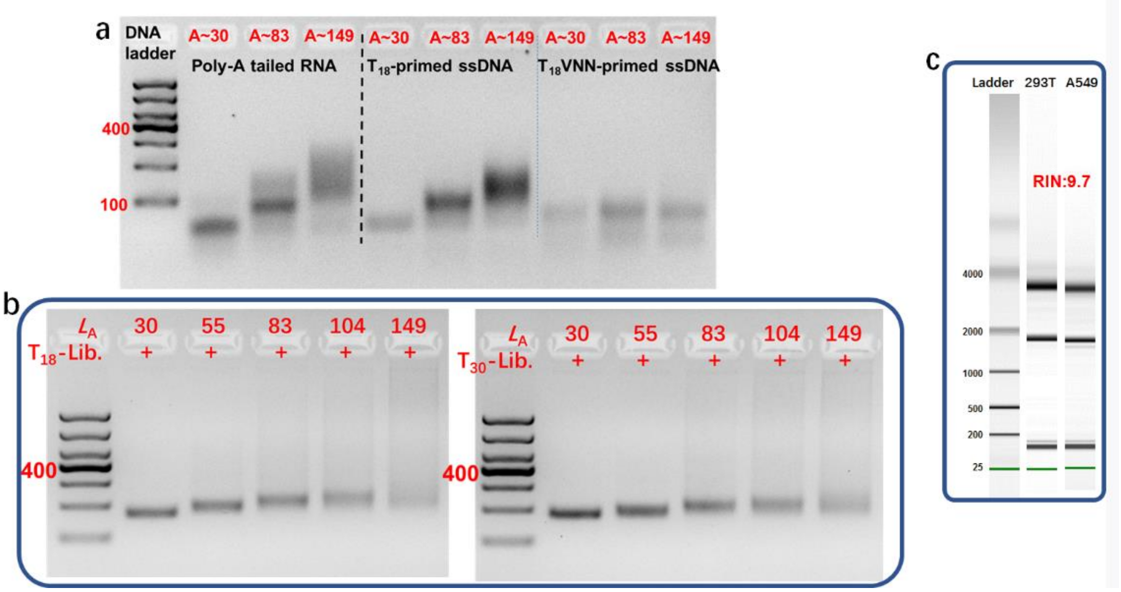


**Supplemental Fig. S1**. Electrophoresis results. **a**) Agarose gel (3%) analysis of spike-in (60nt) with $A_{30}$, $A_{83}$, and $A_{149}$, and its corresponding 1st cDNA upon $T_{18}$ and $T_{18}$VNN priming. **b**) Agarose gel (3%) analysis of sequencing libraries of spike-in with $A_{30}$, $A_{55}$, $A_{83}$, $A_{104}$, and $A_{149}$ based on $T_{18}$ and $T_{30}$ primers. KAPA HiFi 2× master mix is used. **c**) Extracted total RNA from HEK293T and A549 is measured by Agilent 2100 bioanalyzer.

**Fig. S2**

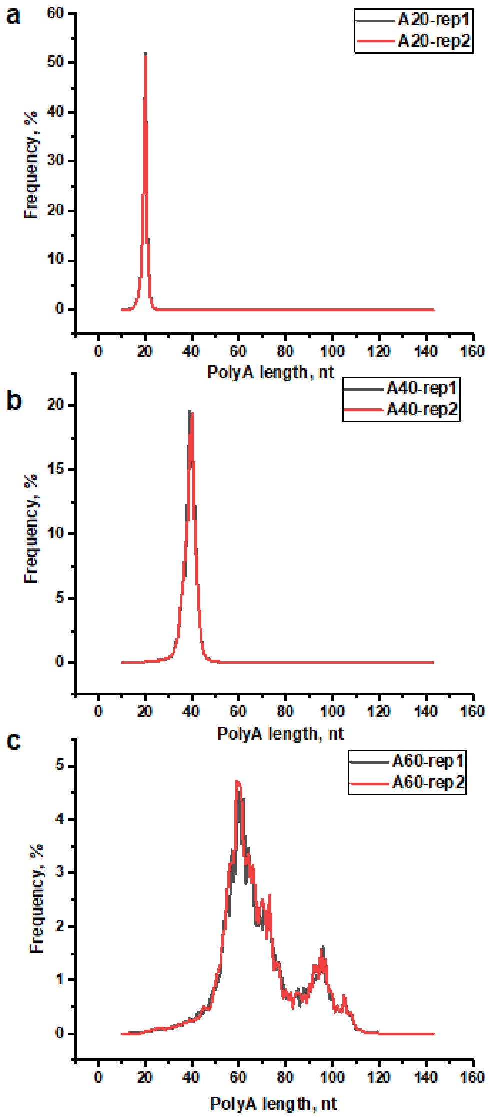


**Supplemental Fig. S2**. Poly(A) length determination of synthesis DNA homopolymer, A20 (**a**), A40 (**b**), A60 (**c**).

**Fig. S3**.

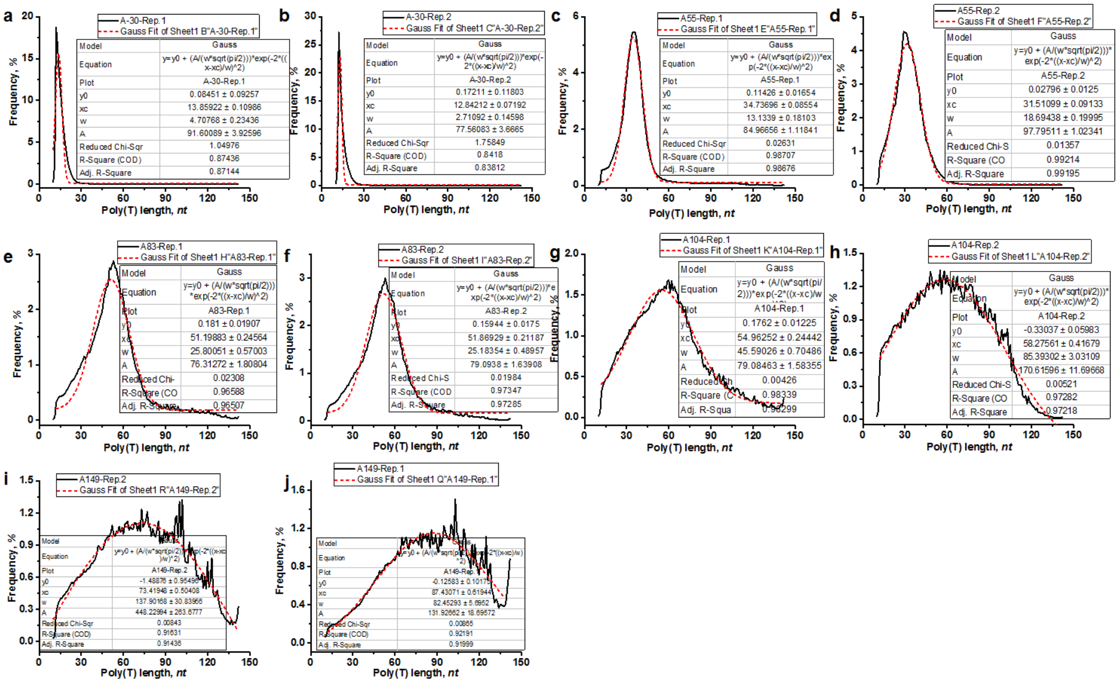


**Supplemental Fig. S3**. CLT-sculpted Poly(T) profile fitting with pseudogaussian function. The spike-ins carrying poly(A) tail of 30 (**a** and **b**), 55 (**c** and **d**), 83 (**e** and **f**), 104 (**g** and **h**), and 149 (**i** and **j**) nt with two replicates are subjected in CLT-seq with $T_{12}$ primer.

**Fig. S4**.

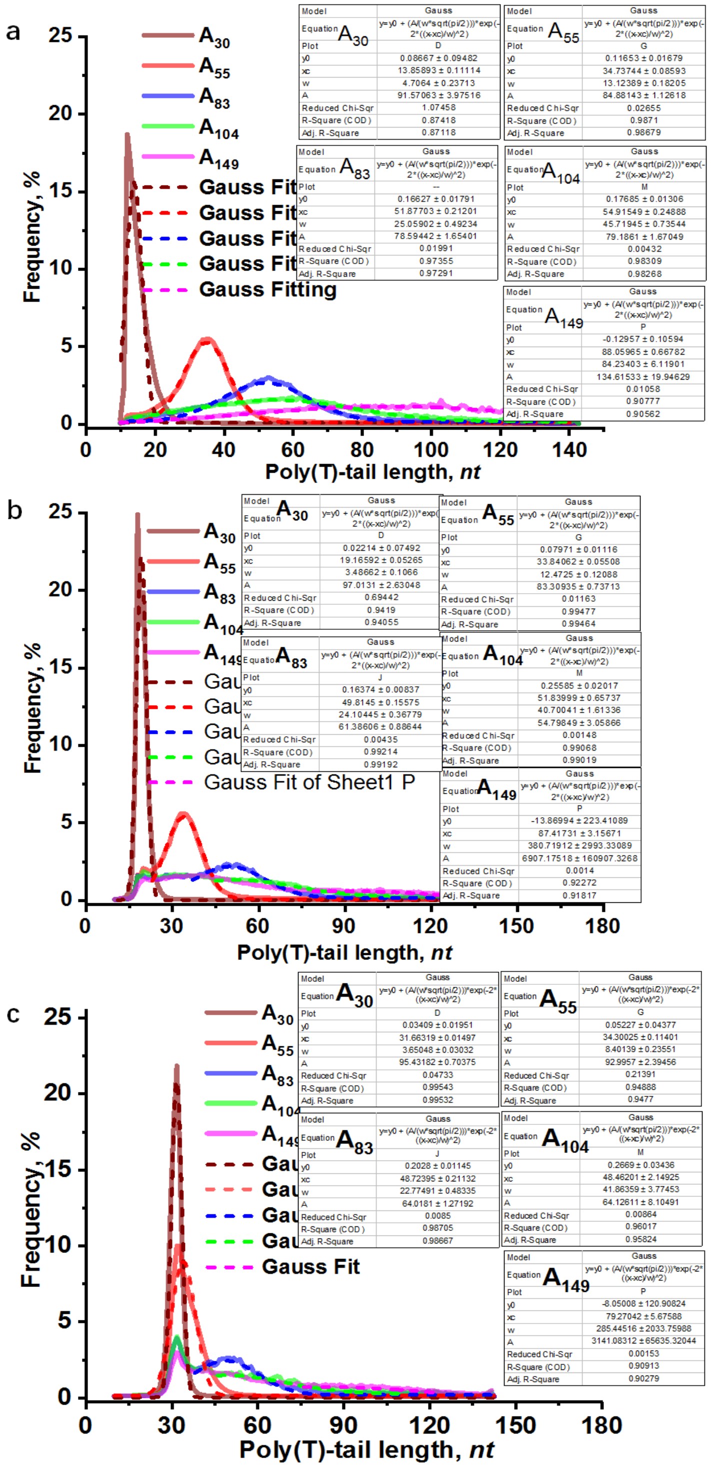


**Supplemental Fig. S4**. Superimposed poly(A)-tail length-dependent CLT-sculpted poly(T) profile with its pseudogaussian fitting curve. The spike-ins carrying poly(A) tail of 30, 55, 83, 104, and 149 nt are subjected in CLT-seq with $T_{12}$ (**a**), $T_{18}$ (**b**), and $T_{30}$ (**c**).

**Fig. S5**.

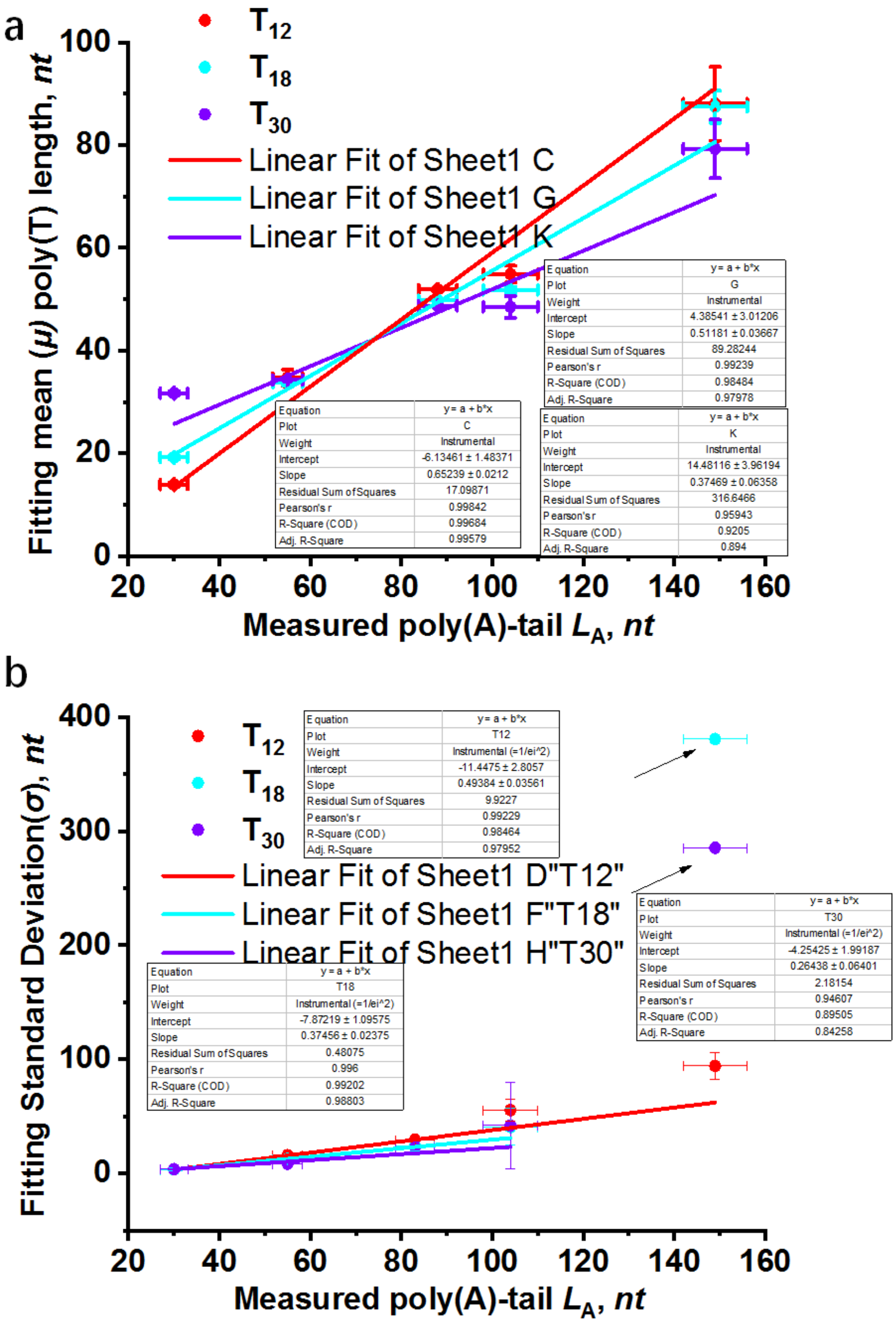


**Supplemental Fig. S5**. Modeling conversion of poly(A) length to poly(T) length underlying CLT mechanism. The spike-ins carrying poly(A) tail of 30, 55, 83, 104, and 149 nt are subjected in CLT-seq. $T_{12}$, $T_{18}$, and $T_{30}$-based regression lines of poly(A) tail length versus fitting poly(T) mean length (**a**) and fitted standard deviation (**b**). Due to a very large error, the standard deviation of spike-in-$A_{149}$-subjected $T_{18}$ and $T_{30}$-based results is not considered for linear regression analysis.

**Fig. S6**.

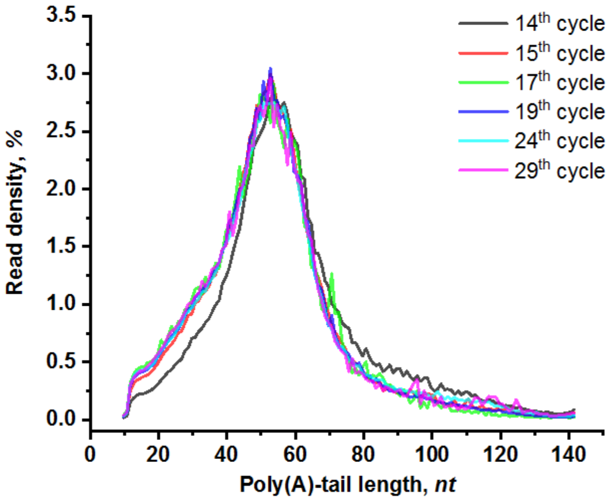


**Supplemental Fig. S6**. PCR bias in poly(T) profiling. The insert profile of each library is overplotted versus counting frequency and GC content.

**Fig. S7**.

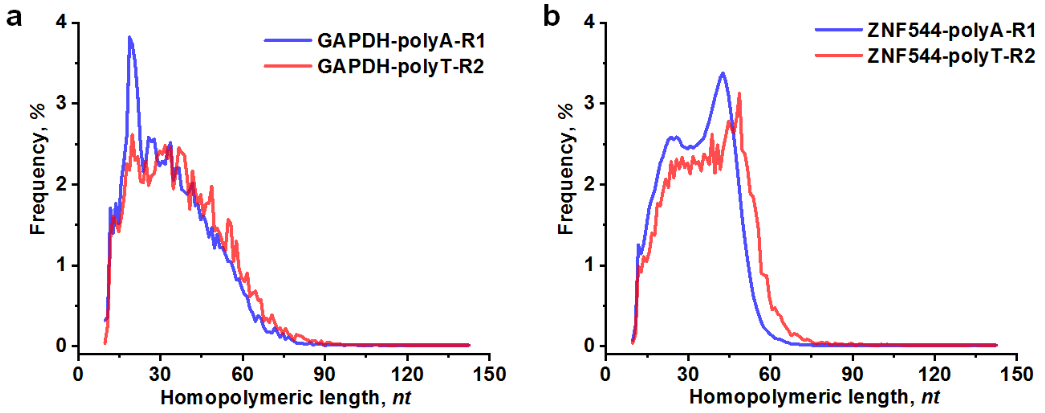


**Supplemental Fig. S7**. Overlapped profile of the R1-outputted poly(A) and the R2-outputted poly(T) from the same pair-end reads for GAPDH (**a**) and ZNF544 (**b**).

**Fig. S8**.

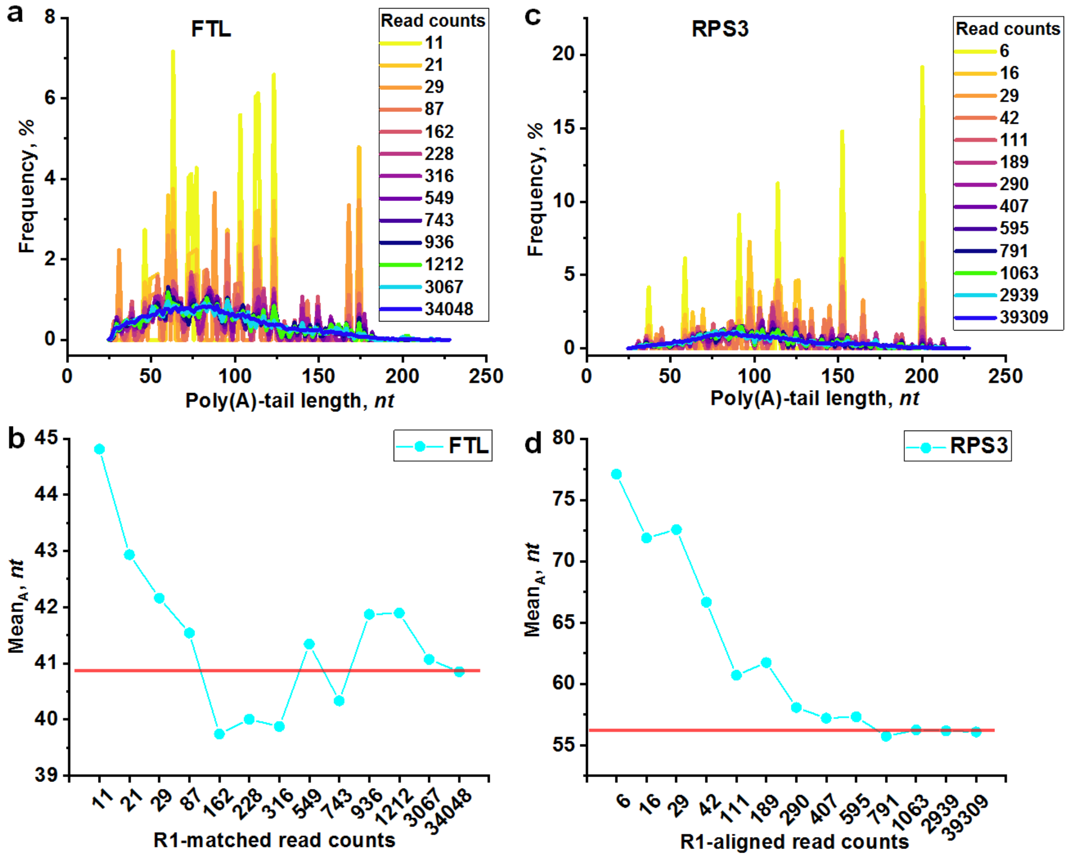


**Supplemental Fig. S8**. Overlapping poly(A)-tail profile with increasing sequencing depth for (**a**) FTL and (**c**) RPS3. The mean poly(A)-tail length plotted for (**b**) FTL and (**d**) RPS3 as sequencing depth increases.

**Fig. S9**.

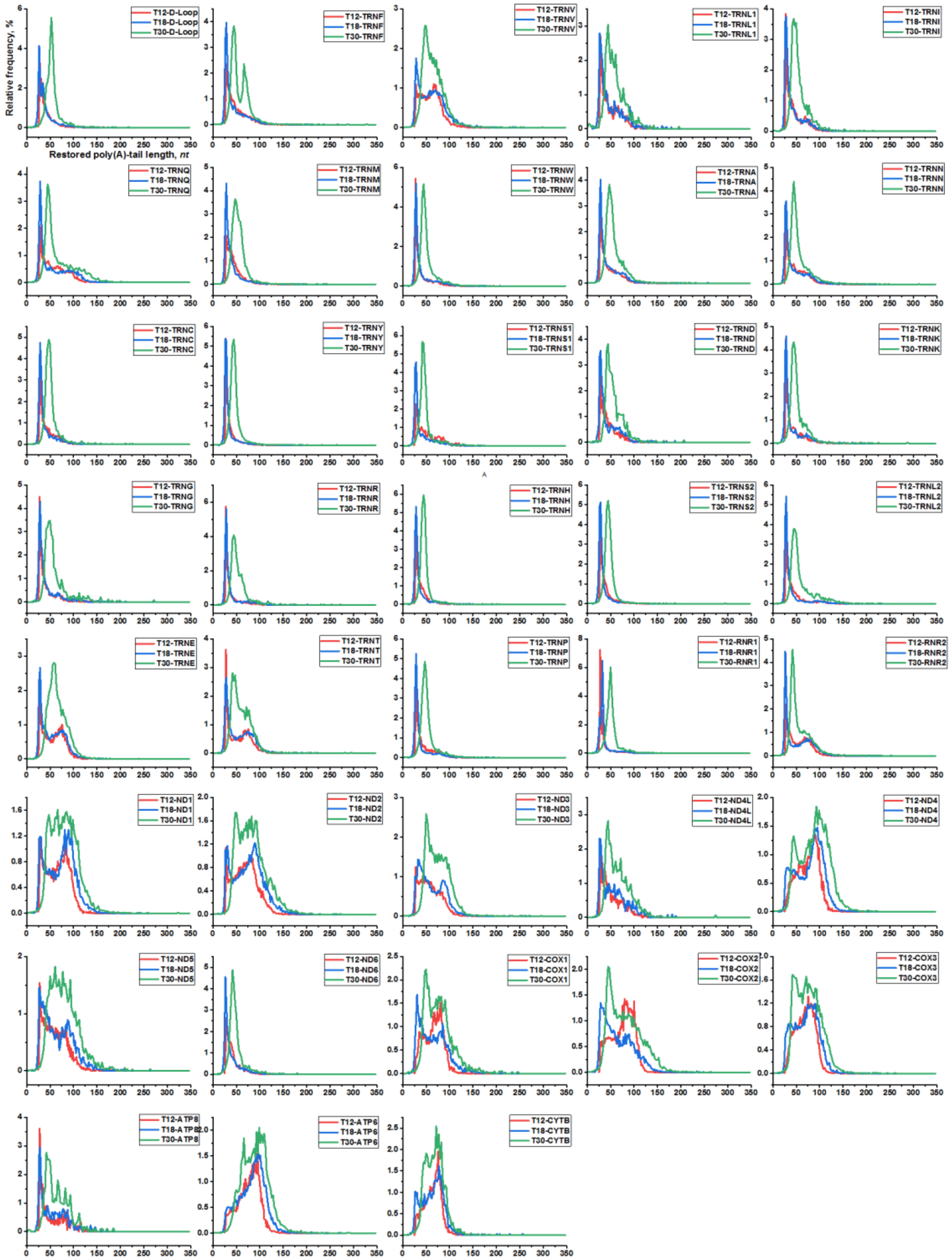


**Supplemental Fig. S9**. The mitochondrial transcript's overlapped poly(A)-tail profile in $T_{12}$, $T_{18}$, and $T_{30}$ primed CLT-seq (13 mRNAs and 25 ncRNAs).

**Fig. S10**.

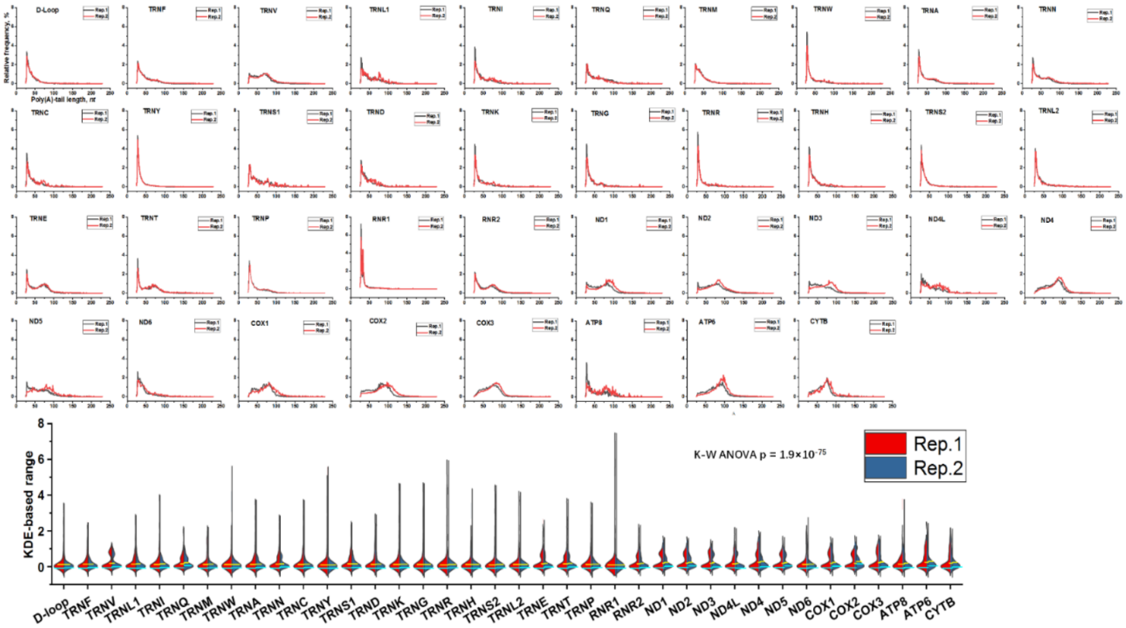


**Supplemental Fig. S10**. Poly(A) tail profile of mitochondrial transcripts in HEK293T. The two replicates superimposed profiles tally counting frequency for each position with single-based resolution. All profiles are also analyzed by the KDE method, and the K-W ANOVA test is measured for pairwise comparison.

**Fig. S11**.

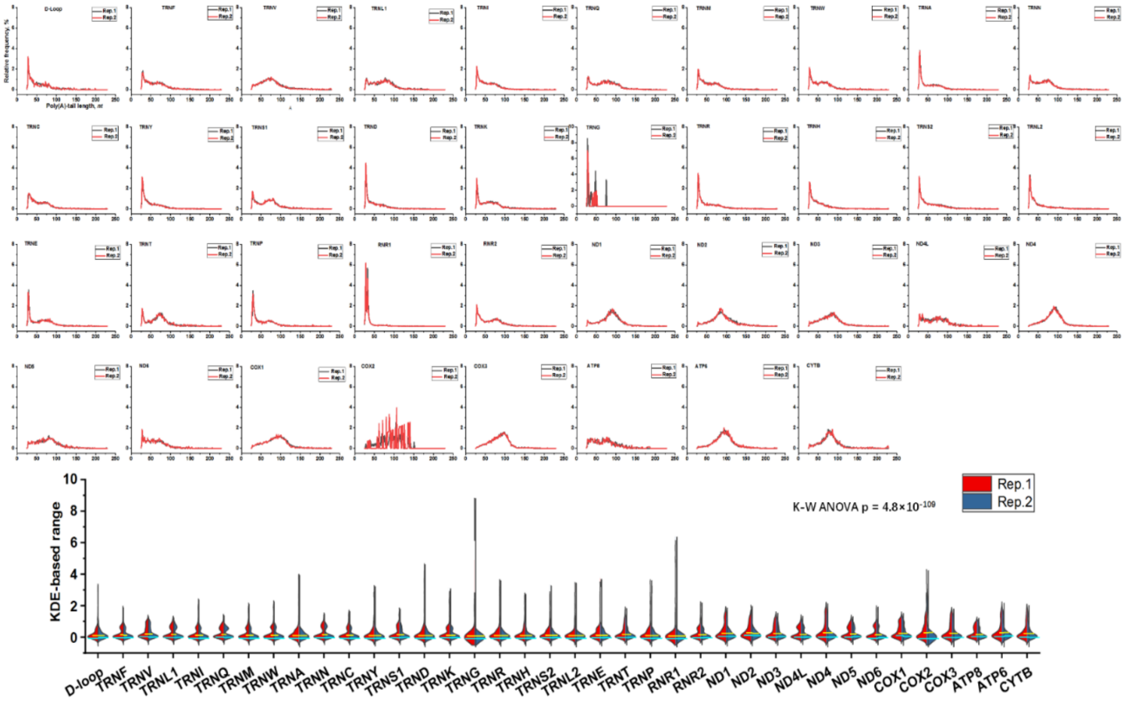


**Supplemental Fig. S11**. Poly(A) tail profile of mitochondrial transcripts in A549. The two-replicates superimposed profiles tally counting frequency for each position with single-based resolution. All profiles are also analyzed by KDE method, and the K-W ANOVA test is measured for pairwise comparison.

**Fig. S12**.

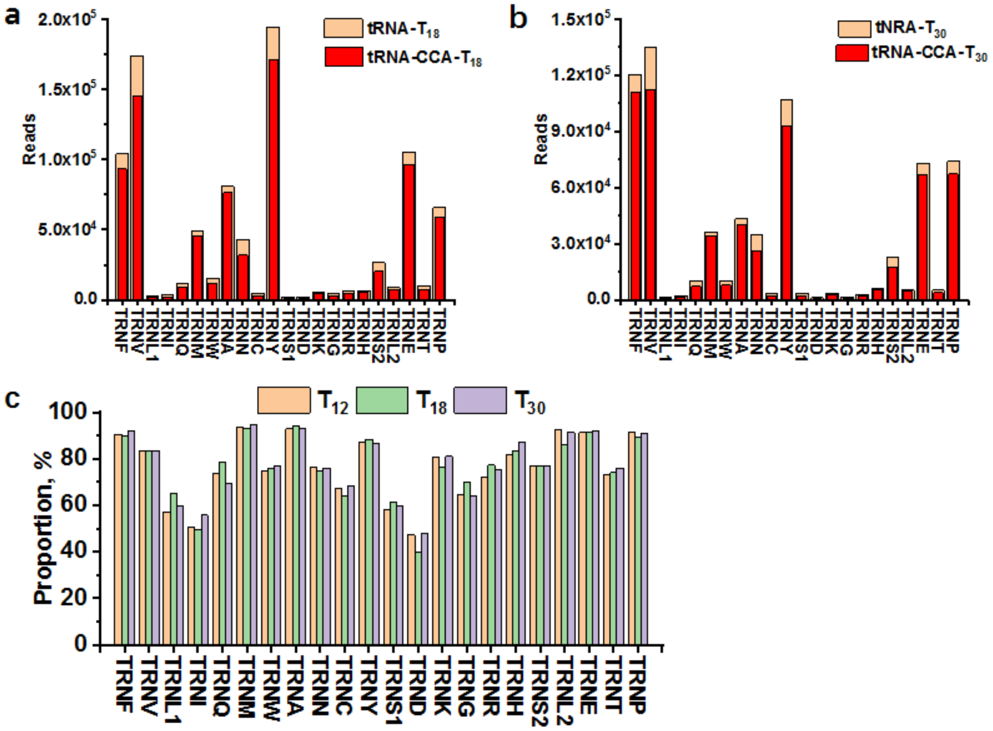


**Supplemental Fig. S12**. The counting reads of CCA marked tRNA and poly(A)-tailed tRNA in all mitochondrial tRNAs of HEK293T upon $T_{18}$ (**a**) and $T_{30}$ (**b**) primed CLT-seq. In addition, the proportion of CCA marked tRNA for poly(A)-tailed tRNA in all tRNAs across $T_{12}$, $T_{18}$, and $T_{30}$ datasets (**c**).

**Fig. S13**.

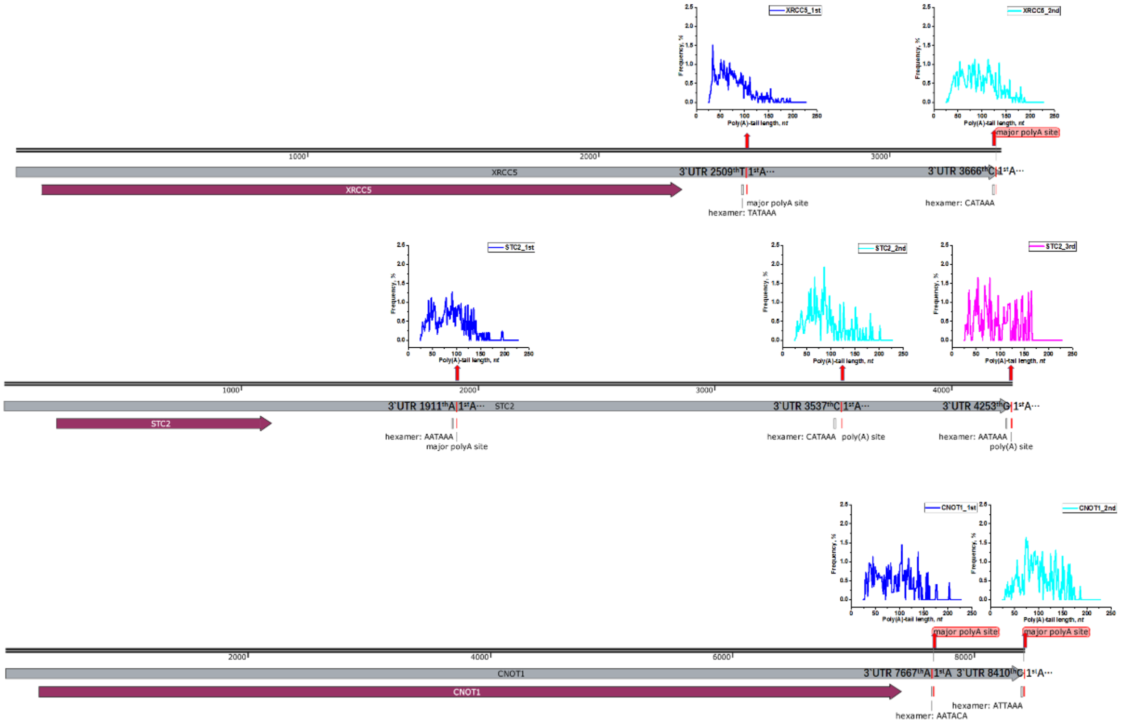


**Supplemental Fig. S13**. Poly(A)-tail profile of isoforms originating from different APA sites in three genes, XRCC5 (up), STC2 (middle), and CNOT1 (down).

**Fig. S14**

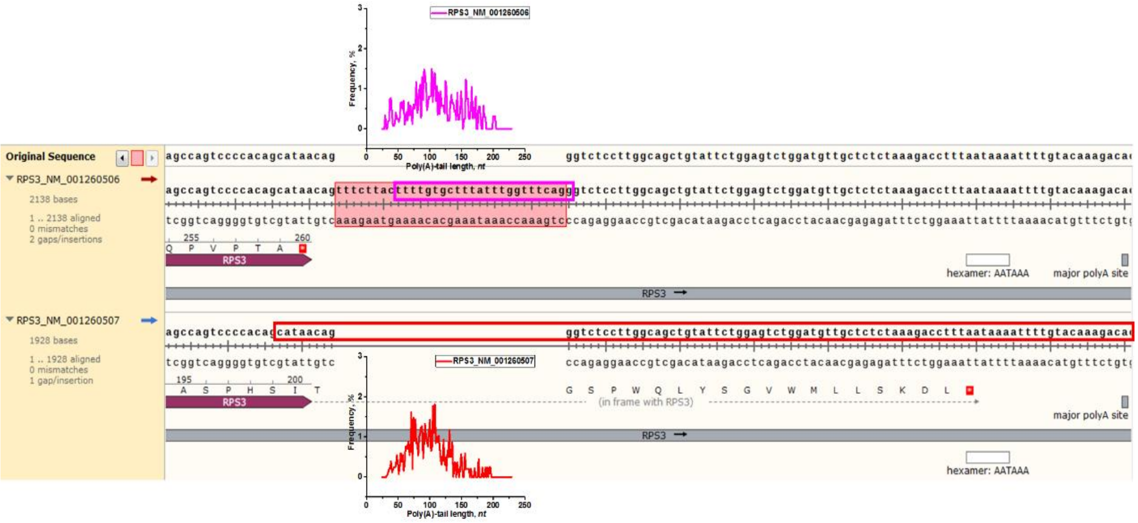


**Supplemental Fig. S14**. Poly(A)-tail profile of isoforms originating from cross-intron spliced isoforms in RPS3 gene.
